# Short-Read Sequencing Benchmarking with Donor-Specific Assemblies

**DOI:** 10.64898/2026.06.24.734333

**Authors:** Sean R. McGee, Joshua D. Smith, Christian D. Frazar, Erica Ryke, Mitchell R. Vollger, Youngjun Kwon, James T. Bennett, Evan E. Eichler, Andrew B. Stergachis, Chia-Lin Wei

## Abstract

**Background:** High-throughput short-read sequencing has become a core technology for genomics, but the rapid expansion of available platforms has made it increasingly important to benchmark them under standardized conditions. A major challenge is that conventional reference-based comparisons confound true sequencing errors with inherited variation and reference bias, making it difficult to isolate platform-intrinsic performance.

**Results:** We benchmarked nine short-read chemistries across seven DNA sequencers using two highly characterized benchmark samples, HG002 and COLO829BL, together with donor-specific assemblies to measure sequencing errors against sample-matched genomic references. This strategy separated authentic platform errors from biological divergence and revealed substantial differences in substitution, indel, read-position, and sequence-context error profiles. Element AVITI UltraQ and Roche SBX-D showed the lowest substitution error rates, whereas Ultima and Roche chemistries exhibited the strongest indel-associated biases. We also found pronounced platform-specific effects in low-complexity regions and trinucleotide contexts, including homopolymer-associated errors and context-dependent substitution skews that are directly relevant to rare-variant detection. In addition, we show that donor-specific references are essential for unbiased base-quality recalibration because they minimize reference bias and more faithfully support cross-platform comparison and low-frequency variant-calling thresholds.

**Conclusions:** Donor-specific assembly-based benchmarking provides a robust framework for measuring true short-read sequencing errors and comparing platforms on a common, sample-matched basis. Our results establish a comprehensive reference for the community and show that authentic error profiles can guide platform selection, quality filtering, and improved detection of rare somatic variation.

## Background

Advances in high-throughput short-read sequencing have transformed genomics into a foundational technology for molecular analysis across research and clinical settings [1–3], enabling applications ranging from population-scale whole genome sequencing [4,5] to single-cell and spatially resolved functional genomics [6]. Over the past decade, improvements in throughput, data quality, and assay design have substantially broadened the reach and impact of mainstream platforms like Illumina HiSeq and NovaSeq X [2,7–11]. At the same time, the field has seen a rapid influx of new sequencing chemistries and instrument systems, many of which are designed to deliver high-throughput, accurate, and cost-effective data while expanding versatility across diverse experimental capabilities [2,8,12–14]. This continued diversification has accelerated the use of short-read sequencing in both discovery and translational applications, where performance is judged not only by accuracy but also by read length, output, run time, scalability, and workflow flexibility. These capabilities are now being deployed to interrogate somatic mosaicism [15] and other rare, low-allele-fraction variants across diverse human tissues. The ability to perform ultra-deep coverage sequencing with improved chemistries and barcoding strategies has increased sensitivity in detection of low-frequency mutations and informed disease risk, clonal dynamics, and therapeutic response.

The broad adoption of genome sequencing has been propelled by the drastic reduction in cost [16], which has in turn enabled greater diversification in sample collection strategies, library preparation, sequencing chemistry, and downstream analysis workflows. As a result, short-read platforms from Illumina, Ultima, MGI, Element, Roche, and PacBio now offer distinct trade-offs in accuracy, error structure, coverage uniformity, throughput, and workflow design, with direct consequences for applications such as somatic mosaicism detection, low-frequency variant calling, and single-cell genomics. The ability to detect somatic variation is central to understanding biological mosaicism, clonal evolution, and disease-associated mutational processes across human tissues [15]. This task is especially challenging at low allele frequencies, where true rare variants must be distinguished from platform- and context-specific sequencing errors that can otherwise obscure or mimic real signal. In this context, the Somatic Mosaicism across Human Tissues (SMaHT) Network [15] was established to systematically characterize somatic variation and to define accurate strategies for its detection across diverse tissues and assay types. Achieving these goals requires accurate, scalable, and robust short-read technologies coupled with optimized analytical workflows, since high-quality detection of rare somatic variants depends on resolving very low variant allele fractions with confidence.

Although selective benchmarking studies and platform evaluations have been reported [10], differences in study design, sample composition, and reference standards make unbiased comparisons difficult. As sequencing technologies continue to diversify and evolve rapidly, there remains a need for standardized, unbiased benchmarking frameworks that can compare accuracy, sensitivity, and reproducibility across platforms using robust reference materials and well-defined metrics. Such systematic evaluation is essential for identifying platform-specific strengths and limitations and for guiding technology choice and optimization in both basic and translational genomics.

A central challenge in achieving this goal is that benchmarking against generalized human references can obscure the distinction between technical error and true biological variation. Because any individual genome contains thousands of single-nucleotide variants (SNVs), small insertions and deletions (indels), and structural variants (SVs) relative to the reference [17], observed mismatches may reflect either genuine sequencing error or inherited divergence, making it difficult to isolate platform-intrinsic performance. This limitation is especially problematic in repetitive and low complexity regions, where reference bias can distort estimates of sequencing accuracy and obscure systematic error modes. Donor-specific assemblies (DSAs) address this problem by providing sample-matched reference genomes built from the same individual being sequenced [18]. By aligning reads directly to the underlying donor sequence, DSAs reduce confounding from germline variation and enable more faithful measurement of platform-specific error rates, context-dependent biases, and base-quality behavior.

In this study, we use DSA-based benchmarking to move beyond conventional reference-based comparisons and more directly resolve the authentic error profiles of multiple short-read platforms. This strategy allows us to quantify mismatch rates, indel burden, and sequence-context effects in a way that more accurately reflects the intrinsic performance of each chemistry. As a result, the DSA framework provides a more rigorous basis for identifying platform-specific limitations, comparing technologies on a common footing, and establishing benchmarking metrics that are optimally suited to applications requiring high-confidence detection of rare variants.

## Results

### Overview of sequencing platforms and benchmarking samples

We systematically evaluated nine short-read sequencing chemistries implemented across a variety of commercially available DNA sequencers. These included Illumina sequencing-by-synthesis (SBS) [19] and the newer XLEAP-SBS implementation on NovaSeq X and X Plus; Element AVITI avidity base chemistry (ABC) and its higher-fidelity UltraQ version [12]; PacBio Onso sequencing-by-binding (SBB) [20]; MGI DNBSEQ-T7 combinatorial probe anchor chemistry (cPAS) [21]; Ultima UG100 flow-based SBS and its duplex paired plus-minus sequencing (ppmSeq) mode [13]; and Roche Axelios sequencing-by-expansion duplex (SBX-D) technology (Table 1) [14]. Together, these platforms represent a broad and currently relevant collection of commercial short-read sequencing technologies used across genomic research applications. Detailed platform specifications are provided in Supplementary Information and Additional file 1: Table S1. We selected two benchmarking samples to enable accurate and broadly relevant cross-platform assessment. HG002, a HapMap-derived cell line widely used by Genome in a Bottle (GIAB) [22], has been extensively characterized by multiple complementary technologies and supported by high-confidence reference materials, making it an established standard for evaluating short-read accuracy. COLO829BL [23], the matched normal lymphoblastoid cell line paired with the COLO829 melanoma cancer line, was included as a SMaHT benchmark resource [24] because it offers a cancer-relevant genome for assessing platform-specific error profiles in a second biologically distinct background. Importantly, both samples have highly resolved, near telomere-to-telomere genome coverage [25], which allow sequencing reads to be compared against sample-specific genomic contexts rather than a generic reference alone. Together, these two samples provide complementary benchmarking datasets that support comparison of error profiles across platforms while maximizing relevance to both germline and somatic sequencing applications.

**Table 1.**
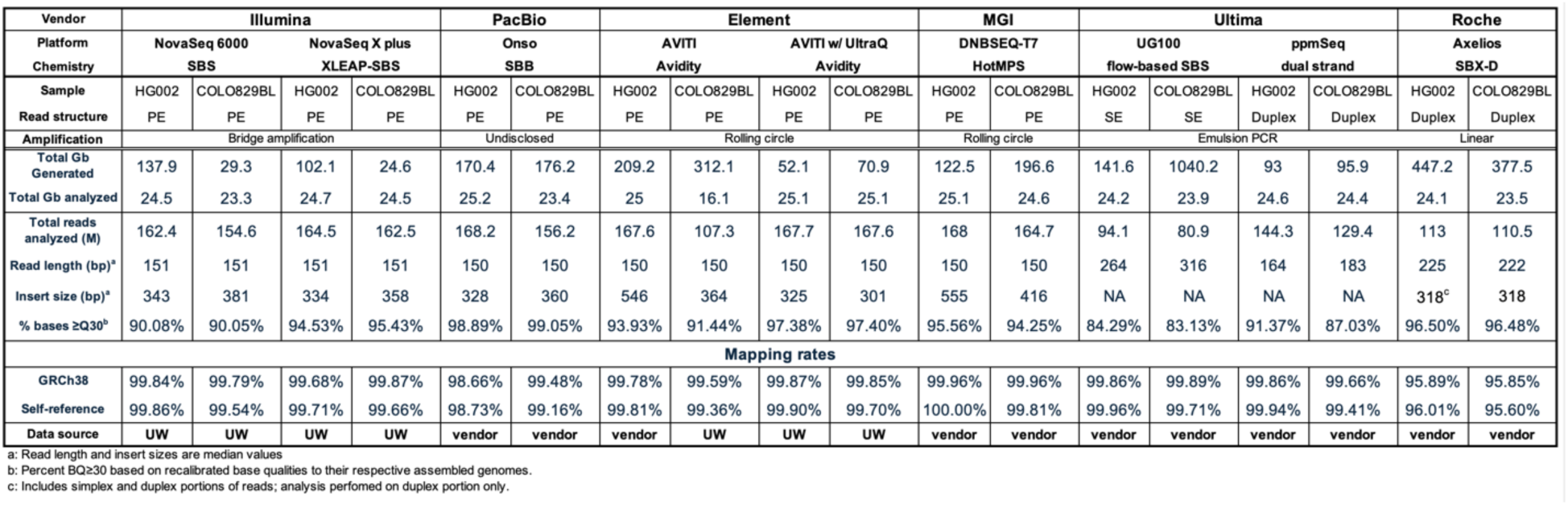
Summary of Data Generated and Analyzed for Each Sequencing Platform.

To minimize batch effects, COLO829BL genomic DNA was extracted in a single batch and then split for library preparation across the different platform-specific workflows. For COLO829BL, all sequencing datasets were generated in-house from the same DNA batch; for HG002, we generated data on the Illumina NovaSeq 6000, NovaSeq X Plus, and Element AVITI UltraQ platforms in-house, and incorporated publicly available datasets from the remaining vendors. In total, we analyzed 18 datasets spanning the nine chemistries evaluated in this study. The total data yield, read counts, read-length distributions, and platform-reported base-quality metrics are summarized in Table 1. Across platforms, raw data generation ranged from 29 Gb to 1040 Gb. Notably, all platforms except Roche SBX-D report conventional Phred-based quality scores, enabling direct cross-platform comparison. Roche SBX-D instead uses a specialized quality assignment scheme, in which all bases in simplex regions of reads are assigned a base quality (BQ) 22, duplex-concordant bases BQ 39, duplex-discordant bases BQ 5, and nucleotides within homopolymer (HP) tracts containing at least one discordant base BQ 25. This scheme explicitly flags positions that are concordant or discordant between the forward and reverse strands during duplex sequencing.

As shown in Fig. 1, platform-reported base quality across read position varied substantially between sequencing systems, even though most bases exceeded Q30. PacBio Onso and Element AVITI with UltraQ consistently produced the highest median base quality (Fig. 1A) and the most right-shifted cumulative minimal-BQ distribution (Fig. 1B). The early-cycle dip seen in both AVITI chemistries reflects platform-specific quality-score modeling, which uses separate training for early and late cycles and applies dark cycling to mitigate end-repair–associated errors. In contrast, Ultima UG100 consistently exhibited the lowest overall reported base qualities, whereas the ppmSeq implementation markedly improved base-quality scores relative to UG100 and approached those observed for Roche SBX-D, consistent with the expected effects of duplex consensus sequencing. MGI DNBSEQ-T7 was the only platform-sample system to show a difference in base quality across samples, with higher reported base quality in COLO829BL than in HG002, likely because the HG002 dataset was generated with an earlier version of the cPAS chemistry. Together, these differences likely reflect both sample-dependent sequence context and chemistry-specific quality modeling.

**Fig. 1.**
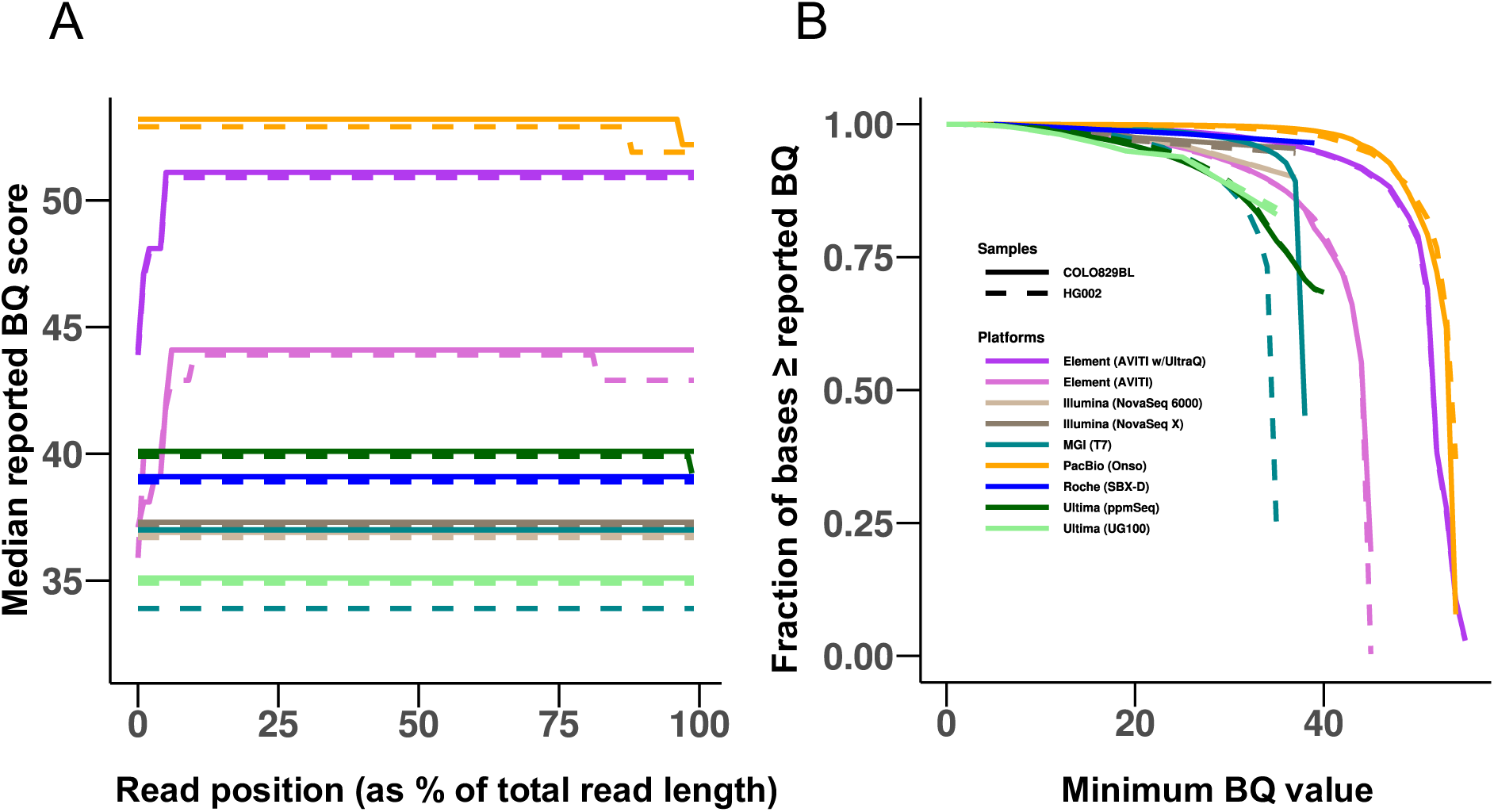
Base-quality score distributions of all nucleotides in sequenced reads for each set of sample FASTQs generated by the nine platforms in this study. **A** The median reported base quality score along the length of the read. Median read lengths in this study range from 150 bp to 316 bp so results are shown as fractional position along the read. **B** Reported-base-quality score distribution for each sample’s analyzed dataset. In panel **A**, median reported BQ values for the COLO829BL and HG002 sample datasets are offset slightly up and down, respectively, from their integer values to avoid overlap.

### Reference-based assessment of substitution and indel errors

Total sequenced data were uniformly downsampled to 24 Gb for error-rate estimation, based on a pilot analysis showing that sequencing error rates could be robustly estimated from 3–54 Gb across a more than 16-fold range in input data (Additional file1: Table S2). We first performed accuracy analyses from the downsampled datasets through alignment to the GRCh38 reference genome. Alignment rates were uniformly high (>95%) across platforms and samples, approaching saturation in most cases, indicating the vast majority of sequence bases can be confidently mapped for accuracy assessment (Table 1). Sequencing accuracy was quantified by calculating mismatch rates for both single-nucleotide substitutions and insertion-deletion events, expressed as the numbers of mismatched events per 1,000 mapped bases (Methods). In these metrics, a mismatch is defined as any position where a nucleotide substitution, or an inserted or deleted segment, differs from the corresponding reference sequence. The single-nucleotide mismatch rate (SNMR) is therefore the number of single nucleotide substitutions per 1,000 mapped bases, whereas the indel mismatch rate (IDMR) is the number of insertion or deletion events per 1,000 mapped bases. To estimate the contribution of genuine germline variation in the reference genome-based error analysis, we simulated an idealized short-read dataset from the COLO829BL genome assembly and mapped it back to GRCh38 under the same alignment conditions (Methods). This simulated alignment yielded an SNMR of 1.6, which serves as a reference point for interpreting the experimental datasets (Fig. 2A, Additional file 1: Table S3). Across platforms, error profiles were closely comparable between HG002 and COLO829BL, and SNMRs spanned a roughly two-fold range from 2.7 to 5.6. Modest sample-dependent shifts were evident, indicating that residual mismatch rates reflect both platform-specific error and sample- and sequence-context effects. PacBio Onso was lowest among the platforms and comparable to Ultima ppmSeq, a duplex-platform, whereas Illumina NovaSeq 6000 and standard Element AVITI showed the highest SNMRs. MGI T7, and NovaSeq X Plus showed broadly similar SNMRs. Application of UltraQ chemistry to Element AVITI substantially reduced SNMRs for both HG002 and COLO829BL, bringing them to levels comparable to those observed with PacBio Onso. This improvement is consistent with UltraQ’s removal of damaged fragments and dark cycling through early cycles prone to end-repair-associated errors. Likewise, the ppmSeq strategy significantly reduced substitution errors in Ultima flow-based sequencing by generating duplex consensus from both strands. Because this analysis was designed to assess primary data without BQ filtering, Roche SBX-D was excluded from this and the following primary data analyses, since, according to the manufacturer’s definition, duplex base-level accuracy requires application of a base-quality threshold.

**Fig. 2.**
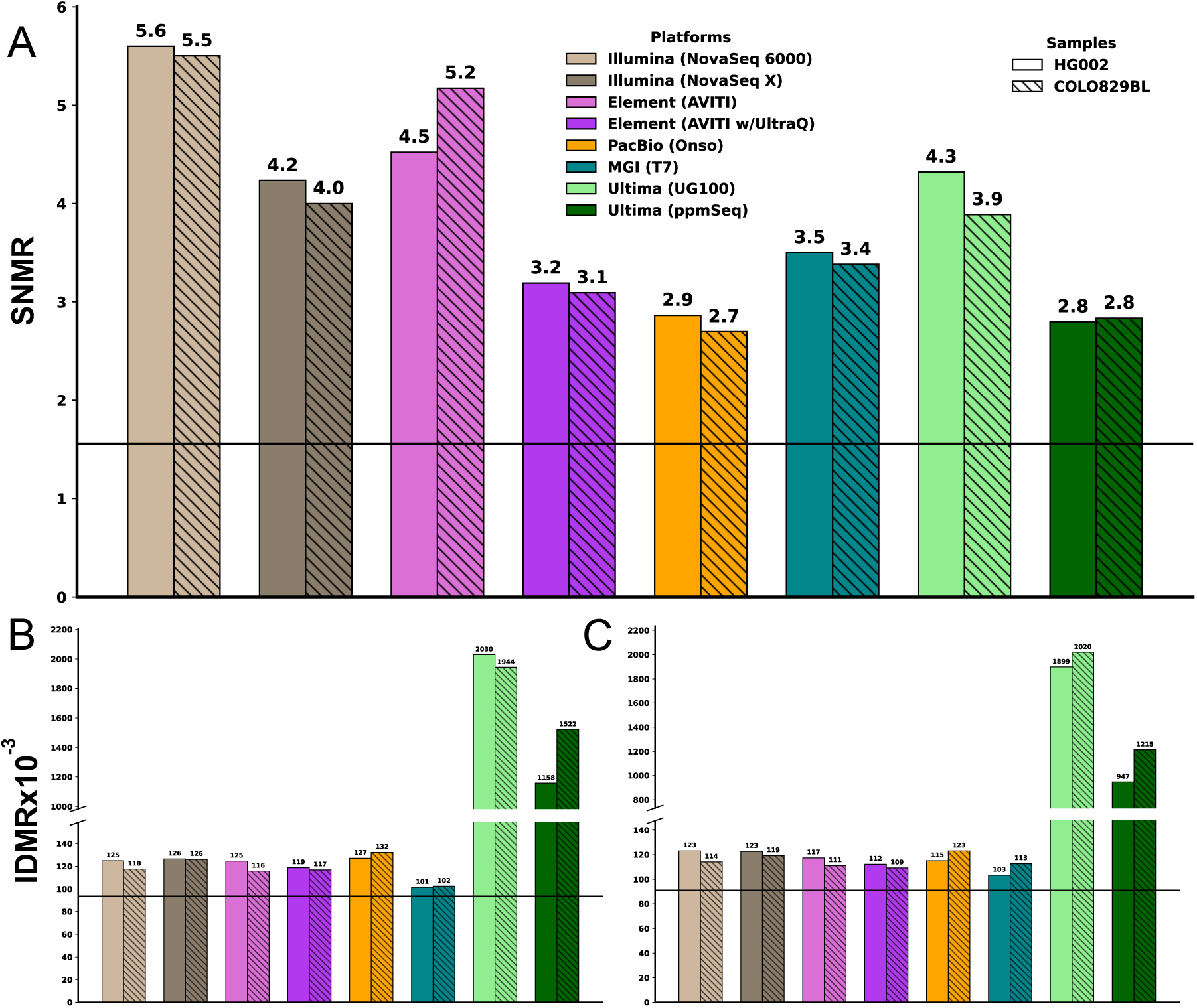
Mismatch rates across sequencing platforms when reads are mapped to GRCh38. **A** Single-nucleotide mismatch rates (SNMR), **B** insertion, and **C** deletion mismatch rates (IDMRs) for COLO829BL and HG002, evaluated across eight sequencing platforms following alignment to the GRCh38 reference. Solid lines in panels **A**–**C** indicate baseline mismatch rates (1.57, 93.7, and 91.2 for single-nucleotide, insertion, and deletion mismatches, respectively) derived from simulated 151 bp paired-end reads generated from the COLO829BL DSA reference and aligned to GRCh38 with *bwa-mem2*. Broken y-axes in panels **B** and **C** are used to display the full range of mismatch rates while preserving resolution among platforms with lower error rates.

IDMRs were next evaluated by analyzing insertions and deletions separately, reflecting the distinct molecular and base-calling mechanisms underlying these two error classes (Fig. 2B and C, Additional file 1: Table S4). Overall, the Ultima chemistries exhibited substantially higher IDMRs, with markedly elevated insertion and deletion errors relative to the other technologies, indicating a pronounced vulnerability of these platforms to indel errors. This is likely reflected by their flow-based chemistries which may increase susceptibility to small shifts in incorporation or phasing that manifest as short insertions or deletions. In particular, Ultima UG100 and ppmSeq showed the largest overall indel burden, with insertion and deletion rates that remained far above those of the other platforms. By contrast, the remaining platforms showed substantially lower and more balanced insertion and deletion rates across samples, generally clustering around ∼100 indel events per million mapped bases. MGI performed comparatively well for indels, with lower rates than the other chemistries. Notably, the COLO829BL and HG002 profiles were broadly similar within each platform, indicating that these indel signatures are predominantly platform-driven rather than sample-specific. Collectively, these results indicate that most platform-specific errors are driven by single-nucleotide substitutions, as indel rates are substantially lower, although higher indel error rates are observed for Ultima chemistries.

### Donor-specific assemblies for measuring true sequencing error profiles

Alignment to GRCh38 provides a useful first estimate of sequencing performance, but the resulting mismatches still reflect a mixture of true platform errors, genuine differences between each sample and the reference genome, and alignment ambiguity in difficult sequence contexts such as low complexity regions. As a result, reference-based error rates can overstate platform error and obscure the intrinsic accuracy of each chemistry. To address this limitation, we next leveraged donor-specific assemblies, which provide sample-matched genomic references and allow reads to be compared against the underlying donor genome itself. This approach enables a more direct assessment of platform-intrinsic substitution and indel errors under a reference framework that minimizes confounding by biological variation. For HG002, we utilized a high-quality diploid donor-specific assembly (DSA) generated by the T2T “Q100” Consortium [26], while for COLO829BL we used a new DSA constructed through the SMaHT Network benchmarking effort using deeply sequenced PacBio HiFi, ultra-long Oxford Nanopore, and Illumina Hi-C data [24,25]. These assemblies provide near telomere-to-telomere representations of each sample’s genome, enabling alignment of short reads directly against their source material to isolate platform-intrinsic sequencing errors from biological variation.

When reads from each platform were aligned to their respective DSAs, SNMRs dropped substantially to a range of ∼0.5 to 3.6 (Fig. 3A), representing a median reduction of ∼2 mismatches per 1000 mapped bases across platforms compared to GRCh38 alignments (Fig. 2A). This consistent improvement confirms that DSA alignment effectively isolates platform-intrinsic sequencing errors by eliminating reference bias from germline variation. Among the eight technologies, PacBio Onso achieves the highest base accuracy (SNMRs 0.5-0.6 across samples), outperforming even duplex methods like Ultima ppmSeq (SNMRs 0.6-0.9), while NovaSeq 6000 showed the highest SNMRs (3.5-3.6), ∼7-fold higher than PacBio Onso. Intermediate value SNMRs (1.6-2.1) were broadly comparable among MGI T7, Ultima UG100 and NovaSeq X. Notably, Element AVITI UltraQ and Ultima’s ppmSeq each delivered 2- to 3-fold improvements over their standard chemistries (standard AVITI 2.4-3.2; UG100 1.8-2.1), underscoring the benefits of damage mitigation and duplex consensus for residual error reduction. Error profiles remained broadly comparable between HG002 and COLO829BL within each platform, reinforcing that DSA-based metrics primarily capture chemistry-specific performance rather than sample effects.

**Fig. 3.**
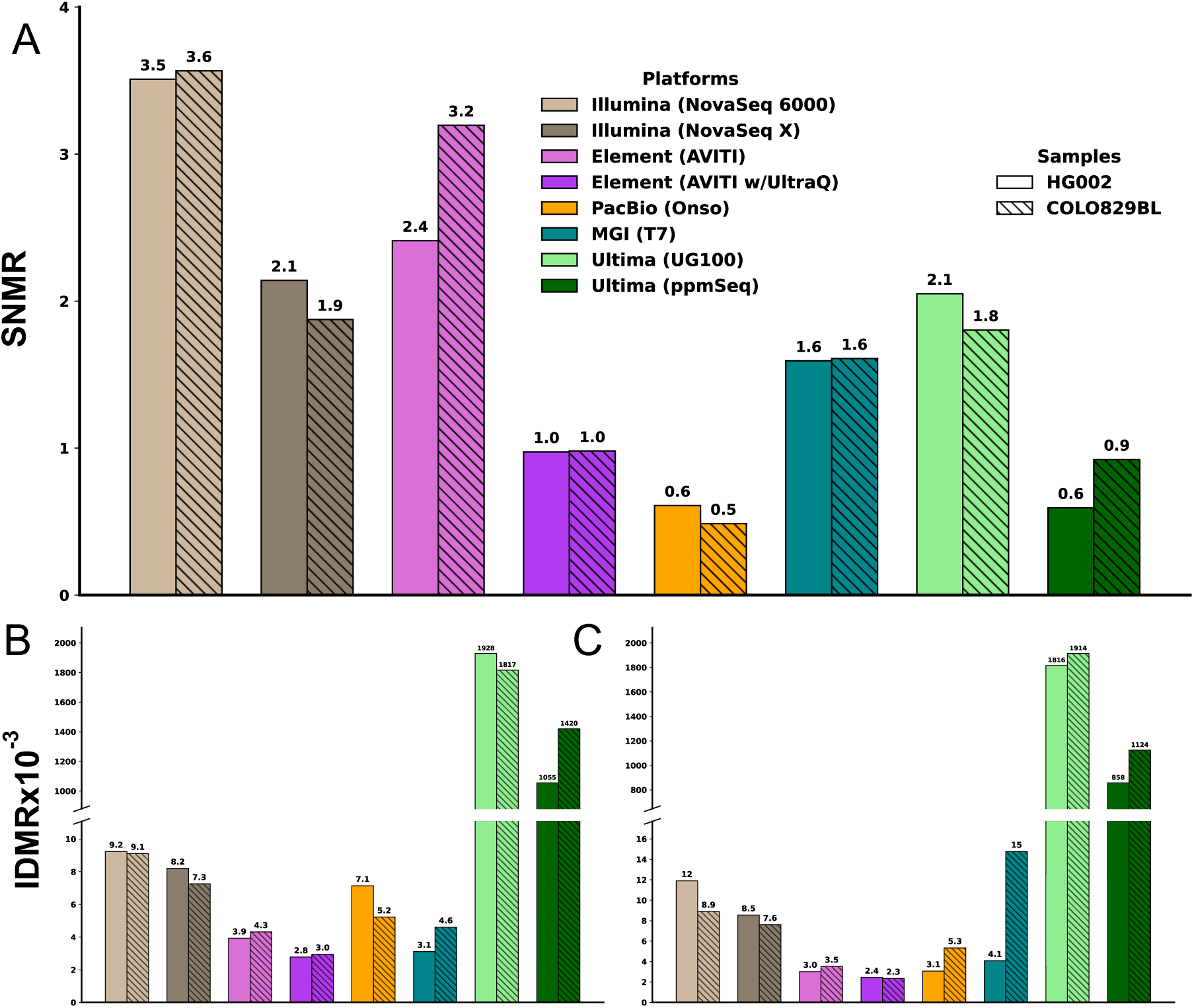
SNMRs and IDMRs for the COLO829BL and HG002 samples across eight sequencing platforms when reads are mapped to their respective donor-specific assemblies (DSAs). **A** single-nucleotide mismatch rates where sequencing reads were aligned to the corresponding custom single-diplotype reference (COLO829BL-DSA for COLO829BL and HG002-DSA for HG002) to minimize reference bias attributable to non-reference alleles. These results represent a measure of the platform-specific base level accuracy under idealized reference conditions tailored to each donor genome. **B** Insertion mismatch rates where, since each sample is being mapped back to its own self-reference, any mismatch could be seen as being strictly an error in the sequencing process. **C** Deletion mismatch rates. Both Ultima technologies have higher indel mismatch rates relative to the other SRS technologies likely due to the specific features of sequencing in those platforms as described in the body of the document. For panels **B** and **C**, a broken y-axis was employed to accommodate the full range of mismatch values observed across platforms, while preserving resolution among platforms exhibiting lower mismatch rates.

DSA-based alignments substantially reduced IDMRs across samples, correcting 93-99% of indel mismatches observed against GRCh38 and yielding residual rates to generally below 10 events per Mb on most chemistries (Fig. 3B and C, Additional file 1: Table S4). This ∼100-fold reduction across all platforms matches the indel contribution from germline variation between the samples and GRCh38, confirming that DSA alignment isolates technical error from biological divergence. Notably, Element AVITI UltraQ consistently achieved the lowest IDMRs (< 3 x 10⁻^6^ for both insertions and deletions), representing the best indel performance among all platforms tested. Ultima’s UG100 and duplex ppmSeq retained markedly higher indel rates (∼1,000–2,000 × 10⁻⁶), approximately 300- to 600-fold above the best performers, likely due to homopolymer length inference errors from signal intensities in their flow-based chemistry. These DSA-resolved error patterns reveal how high-quality DSAs enable direct measurement of true sequencing error rates, revealing performance differences that are confounded when using a generic reference such as GRCh38. By aligning each platform’s reads to DSAs that are effectively sequence-identical to the underlying genomes, residual mismatches predominantly capture platform-intrinsic substitution and indel errors, demonstrating that Element AVITI UltraQ achieves the lowest IDMRs, and duplex flow-based approaches such as Ultima UG100 and ppmSeq exhibit characteristic indel signatures.

### DSA-based recalibration and base quality filtering minimize sequencing errors

Although platforms report base quality scores based on proprietary models and training data, these can be biased by chemistry-specific error modes or training assumptions, necessitating empirical recalibration against alignment outcomes. We therefore performed alignment-based recalibration to derive empirical base qualities for each platform and dataset, enabling direct comparison of error rates as a function of base quality. To test the effects of reference choice on base quality recalibration, we compared recalibrated qualities derived from GRCh38 alignments against those from donor-specific assembly (DSA) alignments. This analysis uncovered marked reference biases in base quality scores, informing optimal quality filtering strategies across platforms. Specifically, alignment-based recalibration of base qualities against GRCh38 systematically underestimates empirical quality scores across all platforms, with recalibrated values consistently falling below the diagonal line (Fig. 4A). By contrast, DSA-based recalibration produces empirical qualities that closely track the manufacturer reported scores, with most platforms aligning near or above the diagonal line (Fig. 4B). This pattern occurs because GRCh38 alignment misclassifies true germline variants as sequencing errors, artificially depressing recalibrated qualities and penalizing platform performance. DSA alignment reduces this reference bias, yielding recalibration curves that more authentically reflect chemistry-intrinsic base-calling accuracy. These results demonstrate that donor-specific references are essential for unbiased base quality recalibration, particularly when comparing diverse sequencing technologies or establishing variant-calling thresholds for low-frequency variants and caution the use of GRCh38-based recalibrations.

**Fig. 4.**
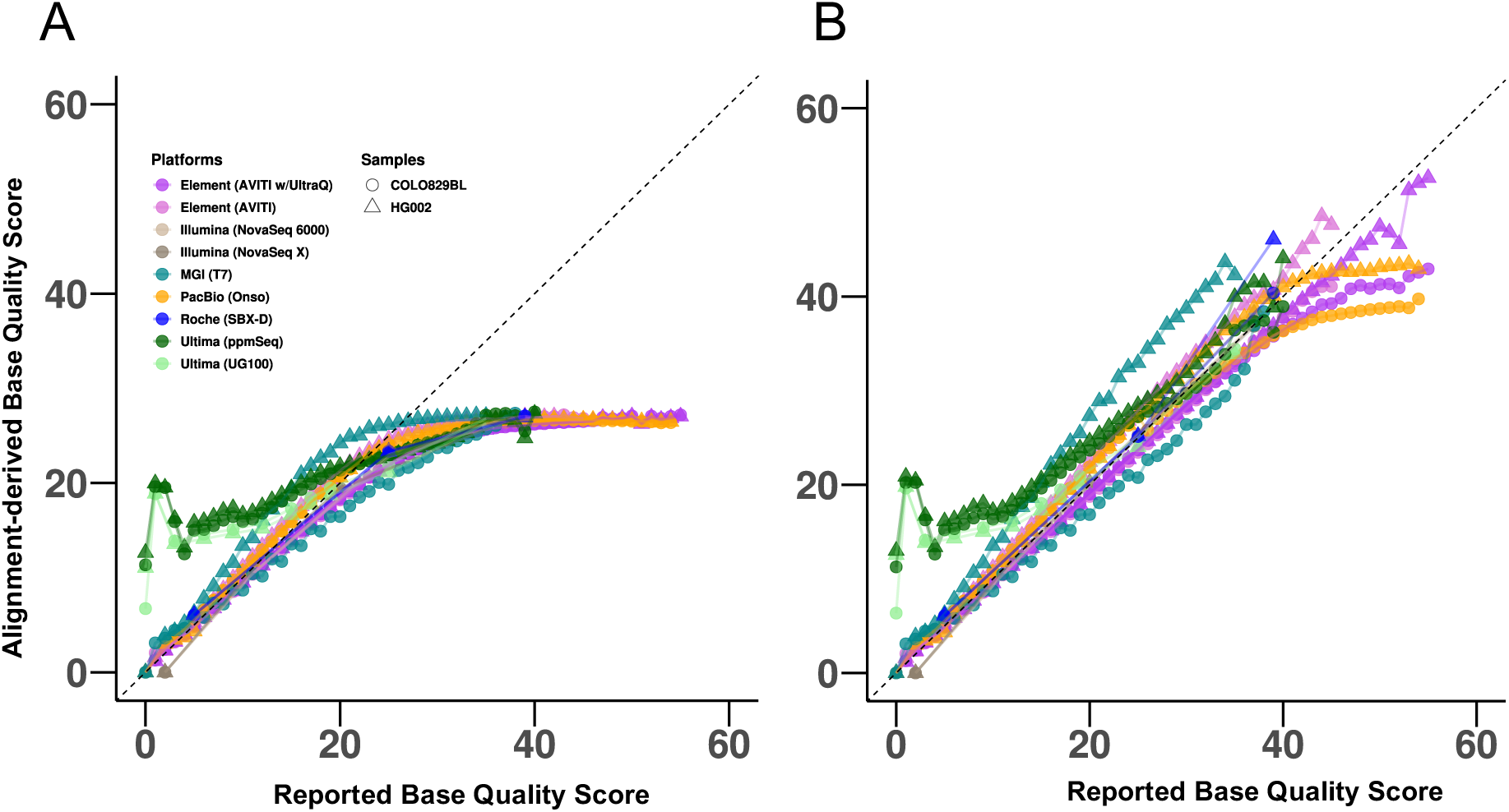
Alignment-based empirical quality scores as a function of the reported base quality score for each platform. **A** COLO829BL and HG002 sample reads aligned to the GRCh38 reference. Recalibrated base-quality scores derived from alignments to a generalized reference genome result in significantly diminished base-qualities. **B** COLO829BL and HG002 sample reads aligned to their respective DSAs illustrates how recalibrations using alignments to a DSA show a much higher correlation to the reported base qualities.

We next examined SNMRs and IDMRs after restricting analysis to bases with DSA-recalibrated BQ ≥ 30, providing a standardized high-confidence subset for cross-platform comparison (Fig. 5). This threshold excluded only ∼1% for PacBio Onso to 17% for Ultima UG100 of total nucleotides across platforms, yet reduced SNMRs by 70%-99% for all chemistries (Fig. 5A, Additional file 1: Table S5). Under this same filtering framework, Roche SBX-D data could also be included in the comparative analyses because its duplex quality scheme is explicitly designed to retain only high-confidence calls after BQ filtering. In particular, a BQ ≥ 30 threshold excludes duplex-discordant bases (BQ 5) and homopolymer-associated nucleotides with discordant signals (BQ 25), while retaining duplex-concordant calls (BQ 39). After filtering, SNMRs converged across platforms to broadly comparable values in the 2.4 × 10^-5^ to 3.5 × 10^-4^ range, including the duplex sequencing platforms Ultima ppmSeq and Roche’s SBX-D, with Roche SBX-D consistently achieving the lowest SNMRs. Ultima UG100, despite substantial filtering, retained a slightly higher SNMR of ∼3 x 10^-4^. Across all platforms, with the exception of Ultima UG100, COLO829BL samples consistently exhibited higher single nucleotide mismatch rates than HG002, with 1.3- to 3.9-fold higher SNMRs. This persistent elevation may reflect culture-derived somatic mutations in the COLO829BL DNA that are not fully represented in the COLO829BL-DSA, or potentially lower assembly quality in the COLO829BL-DSA relative to the T2T-Q100 HG002-DSA. These sample-specific differences highlight the importance of using well-characterized, high-quality reference materials for sequencing benchmarking and underscore a limitation when comparing cell line-derived data against primary reference samples.

**Fig. 5.**
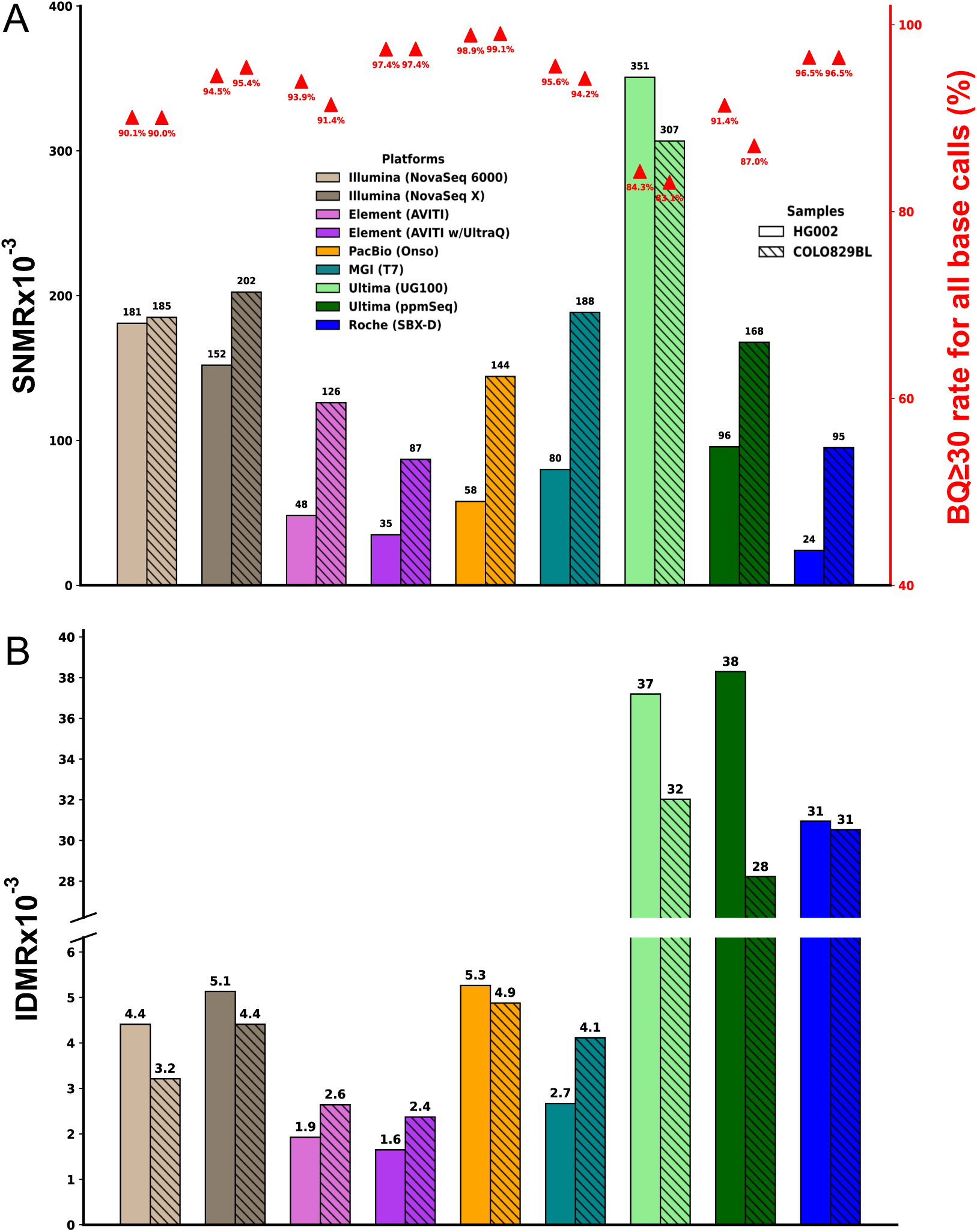
SNMRs and IDMRs for the COLO829BL and HG002 samples across all nine sequencing platforms when reads are mapped to their respective DSAs and the mismatch calculations are restricted to only high-confidence base calls. **A** SNMRs were calculated using only alleles base quality scores ≥30 (BQ ≥ 30). The secondary (right-hand, red) y-axis and labels report the proportion of all base calls meeting the BQ ≥ 30 threshold for each dataset, providing context for the fraction of high-quality bases contributing to the reported SNMRs. **B** IDMRs for insertions for both samples, COLO829BL and HG002, across all nine platforms when mapped to their respective DSA with BQ ≥ 30 filtering of all nucleotides. The rate of high-quality deletions was not considered since there was no consistent way to determine the quality of the deleted sequence (Methods).

We next examined IDMRs for high-confidence insertions (BQ ≥ 30) (Fig. 5B, Additional file 1: Table S5). For most platforms, BQ ≥ 30 filtering provided only modest IDMR improvements, lowering rates to 1.6 - 5.3 events per 10⁶ bases sequenced, because baseline errors were already low. By contrast, Ultima UG100/ppmSeq showed a marked ∼28-57-fold absolute reduction, reaching ∼30 events per 10⁶ bases sequenced, and Roche’s SBX-D exhibited a comparable rate of 31 events per 10⁶ bases sequenced. Compared with SNMRs, indels remained a minor contributor to the overall sequencing error burden for most short-read platforms. Taken together, these results show that DSA-based base recalibration combined with high-quality base filtering removes only a small fraction of bases but eliminates the vast majority of substitution errors, driving SNMRs down to the order of 10^-4^ for nearly all technologies. For indels, most platforms begin with relatively low IDMRs and therefore gain little from filtering, whereas the Ultima chemistries show substantial improvement.

### Platform-specific mismatch biases across read position and sequence context

To further characterize common and platform-specific error behavior, we examined how SNMRs varied by read position and local sequence context using quality-recalibrated reads aligned to the corresponding DSA. This analysis revealed the expected end-of-read deterioration in accuracy, where mismatch enrichment toward later cycles is most pronounced in Element UltraQ, PacBio Onso, and Ultima (Fig. 6A and B), consistent with progressive decline in base-calling performance across the read length. In addition, Illumina and Element AVITI showed spikes of elevated SNMRs at C/G nucleotides near read starts, which may reflect the errors introduced in the end-repair- or library-preparation–associated artifacts rather than true sequencing chemistry alone. These position- and context-dependent effects are important because they can preferentially distort low-frequency variant calls, especially in somatic applications where a small number of systematic errors can mimic rare alleles.

**Fig. 6.**
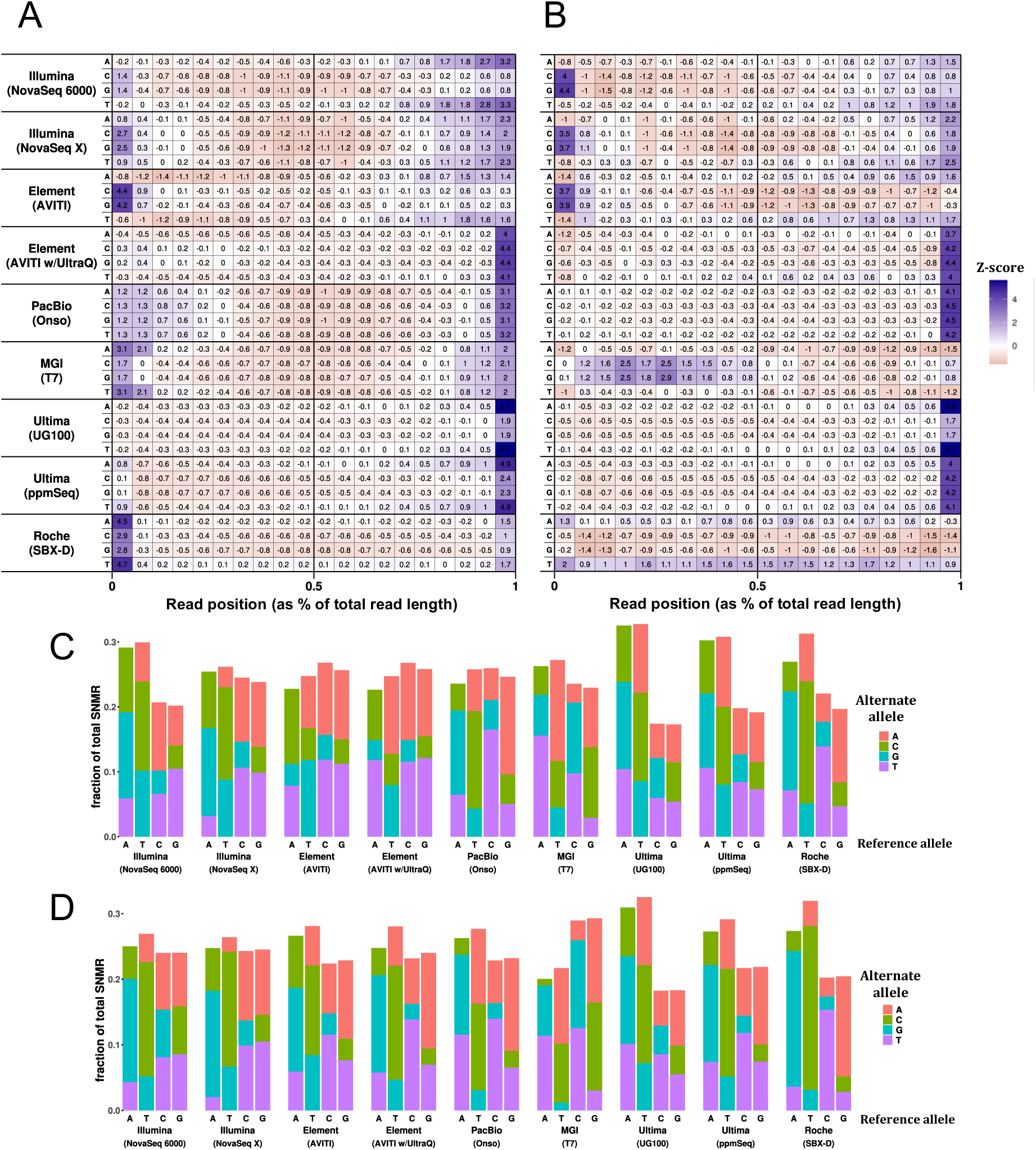
Z-score distributions for single-nucleotide mismatch locations along the read and the fractional contribution of each allele change to the total SNMR for each platform. **A** Z-score distributions for HG002 and **B** COLO829BL, respectively, plotted against nucleotide position along the read after mapping to their respective DSAs. Positions are expressed as the mismatch location relative to total read length due to varying platform read lengths. **C** The fractional contribution of each allele changes for HG002 and **D** COLO829BL when mapped to their DSAs. A notable bias towards reference A- and T-based mismatches is observed across multiple platforms. All analyses are based on highest-confidence mismatches (BQ>=30).

We also observed strong platform-specific substitutional biases. Illumina, Ultima and Roche SBX-D displayed the most pronounced A/T-biased mismatch patterns, whereas MGI T7, particularly in the COLO829BL dataset, showed a distinct G/C-biased signature (Fig. 6C and D, Additional file 1: Table S6), indicating a chemistry-specific error mode that differs from the A/T-skewed patterns seen in several other platforms. As can be seen in the COLO829BL dataset, 64% of the mismatches were at A/T reference bases in Ultima UG-100 and 58% of the mismatches were instead called at C/G reference bases in MGI T7, underscoring substantial context-dependent misincorporation. Among the identities of the substituted bases, Roche SBX-D, Ultima, and Illumina showed the strongest substitution skews overall, with transitional changes dominating the error spectrum. 33% of Illumina and 46% of Roche SBX-D mismatches in COLO829BL were A→G or T→C, representing ∼2.0- to 2.9-fold enrichment over the expected 16% baseline (Fig. 6C and D, Additional file 1: Table S6).

As different sequencing chemistries can generate distinct context-dependent substitution patterns, we determined whether errors were enriched or depleted in specific sequence motifs.

Across platforms, SNM rates showed clear dependence on local trinucleotide context, indicating that mismatch formation is not randomly distributed along the genome. In general, SNMs were enriched in A/T-rich contexts and depleted in GC-rich contexts (Fig. 7A and B), suggesting that sequence composition itself contributes to platform-specific error propensity. This pattern was especially evident for MGI T7, while substitution errors were biased towards to C/G, bases surrounded by C/G were among the least error-prone (Fig. 7C), consistent with a relative depletion of mismatches in NCG/CGN sequence neighborhoods. By contrast, A/T-flanked contexts tended to show higher mismatch burdens across multiple chemistries, supporting the view that sequence-context effects are a major component of residual error after DSA-based recalibration. Notably, the AAA/TTT homopolymer context exhibited the highest SNMRs in the Ultima platforms, consistent with the known challenge of accurately resolving homopolymer length in flow-based sequencing chemistries. Together, these results indicate that platform-specific mismatch profiles are shaped not only by read position but also by local nucleotide context, with important implications for low-frequency variant calling in sequence-complexity-sensitive regions.

**Fig. 7.**
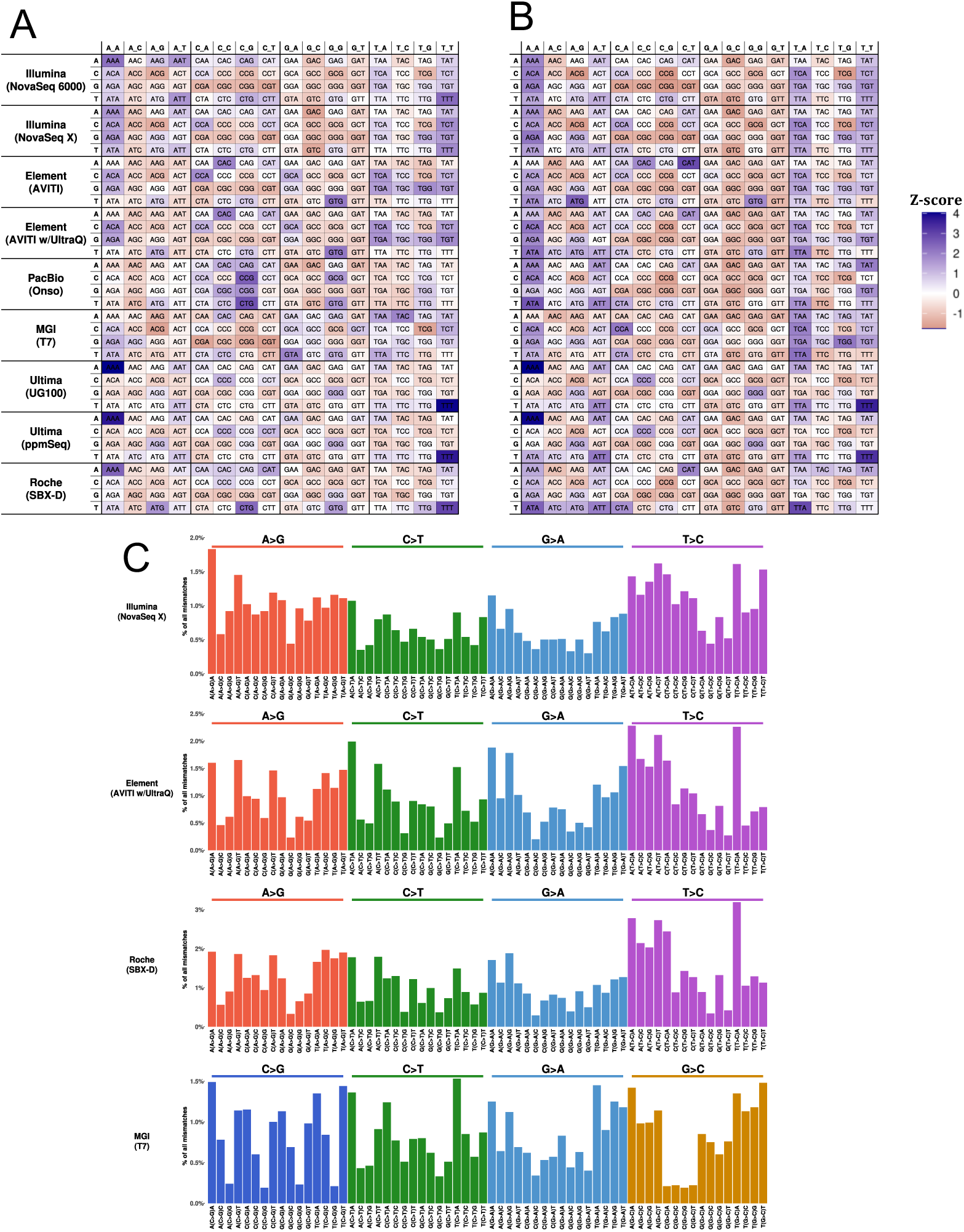
Z-score distributions of the genomic context dependence of single-nucleotide mismatches and the fractional contribution of a specific set of allele changes of interest within the range of genomic contexts. **A** Z-score distribution of HG002 and **B** Z-score distribution of COLO829BL samples mapped to their respective DSAs for each genomic context in which the mismatch occurred. Both plots are based on BQ>=30 single-nucleotide mismatches for each sample mapped to its respective DSA. **C** A set of allele changes for platform/sample combinations of interest detailing the resulting mismatched allele and the specific genomic context in which the mismatch occurred for BQ>=30 single-nucleotide mismatches in the COLO829BL sample mapped to the COLO829BL-DSA. The y-axis is the percentage of total allele changes in each genomic context for all instances for each platform.

### Low-complexity regions contribute disproportionately to platform-specific errors

Given that sequence complexity can strongly shape local error rates, we next asked how much of the substitution and insertion signals are enriched in low-complexity regions (LCRs) such as tandem repeats and homopolymers. Although LCRs account for only a minority of the genome [27], these contexts were known to challenge base calling and alignment and contributed disproportionately to errors affecting both single-nucleotide variant, small indel and structural variation detection [28]. Using the DSA-based error estimates as a cleaner baseline, we directly quantified the extent to which low-complexity and repetitive sequence contribute to sequence errors across different platforms.

In total, LCRs make a higher contribution to the error rate; with the effect being much more evident for single-nucleotide mismatches than for insertions (Fig. 8A and B, Additional file 1: Table S7). Across platforms, SNMs were consistently more highly concentrated in LCRs than in non-LCRs, but the magnitude of this enrichment varied substantially by chemistry. Illumina showed the largest LCR-associated increase in SNM concentrations, whereas Element AVITI, PacBio Onso, and MGI T7 were comparatively similar between LCR and non-LCR regions. By contrast, Ultima UG-100 exhibited the highest concentrations in both sequence classes, with a particularly pronounced SNM elevation in LCRs, while Element AVITI with UltraQ showed the lowest SNM burden overall.

**Fig. 8.**
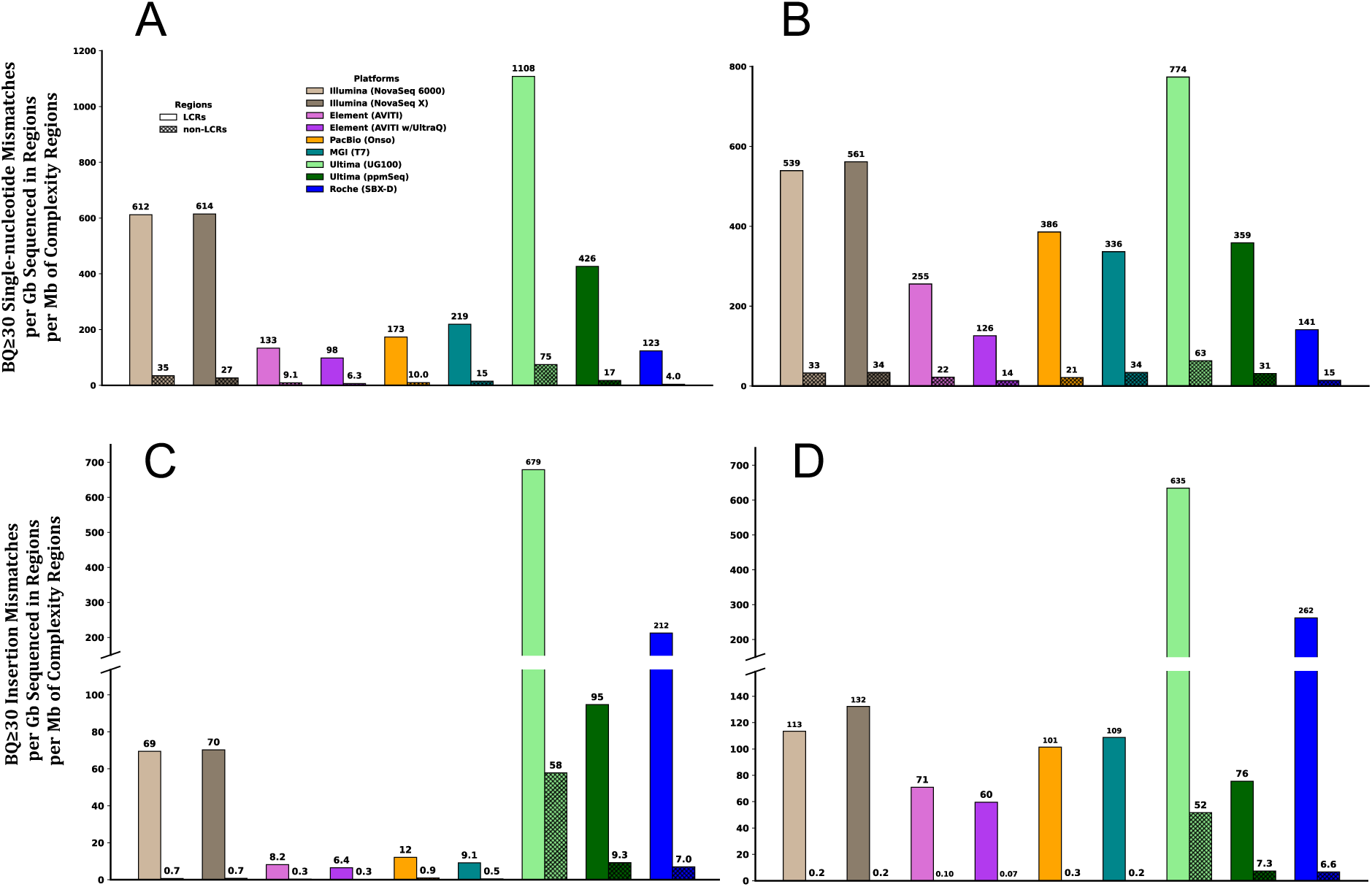
Comparison of the single-nucleotide and insertion mismatch concentrations between low-complexity regions (LCR) and regions outside of these LCRs (non-LCR) when mapped to their respective DSAs. Values are normalized to both the total number of sequenced bases falling within the LCRs as well as to the relative sizes of the LCRs (Methods) **A** HG002 and **B** COLO829BL SNMs. **C** HG002 and **D** COLO829BL insertion mismatches. For panels **C** and **D,** a broken y-axis was employed to accommodate the full range of mismatch values observed across platforms, while preserving resolution among platforms exhibiting lower mismatch rates. LCRs are defined by the GA4GH consortium’s summary of tandem repeat and homopolymer regions in GRCh38 and then these regions were lifted-over to the COLO829BL and HG002 DSAs. Non-LCR refers to all regions of the genomes outside of the LCRs (Methods).

In COLO829BL, the overall relationship between sequence complexity and mismatch burden was broadly concordant with that observed in HG002 (Fig. 8A and C vs. B and D, Additional file 1: Table S7). These patterns indicate that repetitive and compositionally biased regions remain a major source of platform-specific residual error, even after donor-specific calibration.

## Discussion

In this work, we provide a minimally biased benchmarking comparison across the current short-read sequencing landscape, evaluating nine chemistries across seven DNA sequencers on two biologically distinct, highly resolved samples, HG002 and COLO829BL. By pairing all existing short-read platforms with donor-specific assemblies, we separate true sequencing artifacts from germline divergence and residual reference bias, revealing each chemistry’s authentic error profile rather than a mixture of biology and technology. This framework showed that platform performance is highly heterogeneous; Element UltraQ and Roche SBX-D achieved the lowest substitution error rates when filtered for high-quality base calls, Ultima and Roche showed stronger indel burdens, and multiple platforms displayed pronounced read-position, sequence-context, and low-complexity–dependent biases. Beyond ranking technologies, the study also exposes the molecular signatures of error formation, including end-cycle decay, homopolymer sensitivity, and chemistry-specific trinucleotide preferences, all of which are highly relevant for low-frequency variant calling. Collectively, these results establish a high-resolution benchmark for the community and demonstrate that donor-specific assemblies are a powerful way to measure real sequencing error and to guide platform selection, quality filtering, and downstream analysis in both germline and somatic genomics. This is particularly valuable for assessing performance in low-frequency somatic variation, where the distinction between true signal and platform-induced noise becomes increasingly critical. By leveraging a personalized or synthetic “truth set” reference, the limitations of a given sequencing modality can be more accurately quantified, which can facilitate further pipeline refinement to maximize both sensitivity and specificity in variant detection workflows.

While mapping sequencing data back to sample-specific DSAs provides a quantitative framework for evaluating the accuracy of variant allele calls across a range of short-read sequencing platforms, DSAs only improve error estimation if the assembly itself is sufficiently accurate, as low quality and misassembled sequences can amplify platform errors. Here, the HG002 DSA is expected to support a more faithful error assessment because it has been extensively manually polished post-construction [27], whereas the COLO829BL DSA appears to retain more unresolved assembly artifacts [25]. Furthermore, the COLO829BL cells appear to have accrued somatic mutations during culture [29], resulting in higher residual SNMRs. Therefore, a DSA-based benchmarking strategy is most powerful when the assembly is highly contiguous and well validated, and the culture used for DNA extraction has a minimal burden of somatic variants. For that reason, any cross-platform comparisons based on DSAs should be interpreted in light of assembly and DNA quality, especially when comparing datasets from cancer-derived or structurally complex genomes.

Our findings show that base-quality recalibration is highly sensitive to the choice of reference framework, with GRCh38-based approaches consistently under-calling empirical base quality relative to donor-specific assemblies. This underestimation likely reflects the inclusion of genuine germline differences as apparent sequencing errors, which depresses recalibrated quality scores and can lead to overly conservative filtering thresholds. By contrast, DSA-based recalibration more closely recapitulates manufacturer-reported base qualities, suggesting that sample-matched references provide a truer estimate of platform performance and more reliable inputs for downstream analysis. This distinction is especially important for applications in which base quality directly influences variant detection, such as low-frequency somatic mutation calling, mosaicism analysis, and other assays that depend on accurately distinguishing true biological variation from technical noise. More broadly, these results highlight that reference choice can shape not only benchmarking outcomes but also the biological conclusions drawn from sequencing data, particularly when evaluating rare variants near the detection limit.

Although our analyses focused on platform-intrinsic error rates, short-read sequencing now supports a wide spectrum of applications well beyond variant detection, including bulk and single-cell multiomic profiling of genome state, genome perturbation, chromatin accessibility, DNA modification, and spatial or temporal tracking of cellular and tissue dynamics [30–33]. In these settings, accuracy remains essential, but platform choice is increasingly shaped by value rather than accuracy alone, because cost per Gb, turnaround time, throughput, read length, workflow complexity, and assay-specific requirements can be just as important for study design and interpretation. Among the chemistries analyzed, Element UltraQ and PacBio Onso carried the highest cost per Gb, whereas Ultima UG100 was the most cost-efficient, making it particularly attractive for high-depth counting applications. At the same time, the rapid introduction of newer, higher-throughput, lower-cost platforms such as Element Vitari, MGI T7+ and Ultima UG200 continues to shift the performance-cost landscape, underscoring the need for benchmarking frameworks that remain relevant as commercial technologies evolve. Accordingly, benchmarking should inform not only which technology is most accurate, but which is most fit for purpose across diverse biological and translational applications.

In summary, with the continued advances in sequencing technologies and the reduction in the cost of generating high-output, highly accurate short-read data, we are now entering an exciting era in which the knowledge of human genomes can be realized at population scale, creating new opportunities to address unmet needs in human disease research and precision medicine.

## Conclusions

Aligning short-read data to donor-specific assemblies provides a robust framework for quantifying platform-intrinsic sequencing errors and for evaluating the confidence of downstream variant calls. By minimizing confounding factors, like those from germline variation and reference bias, this strategy enables direct comparison of multiple short-read technologies on a standardized, sample-matched genomic baseline and reveals chemistry-specific error profiles that are not apparent from generic reference-based analyses alone. In this study, this approach uncovered substantial differences in substitution, indel, positional, and sequence-context-dependent error patterns across nine different short-read platforms currently available in genomic community, including errors that are particularly relevant to low-frequency variant detection. More broadly, our results show that high-quality donor-specific references can sharpen benchmarking of existing sequencing technologies and provide a practical foundation for platform selection and improved somatic variant discovery in human genomes.

## Methods

### Preparation of HG002 and COLO829BL DNA samples for sequencing data generation

Pure populations of COLO829BL lymphoblastic suspension cells (ATCC cat # CRL-1980, lot # 70022927) were supplied by the SMaHT Network. All cells were grown at 37°C with 5% CO_2_ and extracted in one batch to avoid batch-to-batch variability. DNA was extracted from cell lines and tissues using the Qiagen DNAEasy Blood and Tissue kit (cat #69506). The integrity of the extracted DNA was confirmed using the Agilent Femto Pulse instrument and the DNA yield was confirmed using the Invitrogen Qubit kit and Qubit 4 fluorometer.

### Illumina SBS NovaSeq 6000 and X plus

Starting with a minimum of 750ng of DNA, samples were sheared in a 96-well format using a Covaris R230 focused ultra-sonicator targeting 380bp inserts. The resulting sheared DNA was cleaned with Takara NucleoMag beads to remove sample impurities prior to library construction. Shearing was followed by size selection, and sample prep was performed using the KAPA Hyper Prep kit (KR0961 v1.14). End-repair, A-tailing, and ligation were performed as directed. Two final NucleoMag cleanups were performed after ligation to remove excess adapter dimers from the library. All library construction steps were automated on the Revvity Janus platform. QPCR was performed using the Bio-Rad CFX384 Real-Time System and the KAPA SYBR FAST qPCR Master Mix (079559397001) in order to determine the nanomolarity of the final libraries. Sequencing was carried out on the NovaSeq 6000 and X Plus sequencers. Base calls were generated in real-time on the instrument (RTA v3.4.4 and v4.29.2, respectively) and then demultiplexed, with fastq files produced by bcl convert v4.2.7.

### AVITI with Cloudbreak and UltraQ

For standard AVITI libraries, starting with a minimum of 750ng of DNA, samples were sheared in a Covaris R230 focused ultra-sonicator targeting 380bp inserts. The resulting sheared DNA was cleaned with Takara NucleoMag beads to remove sample impurities prior to library construction. Shearing was followed by size selection, and sample prep was performed using the KAPA Hyper Prep kit (KR0961 v1.14). End-repair, A-tailing, and ligation were performed as directed. Two final NucleoMag cleanups were performed after ligation to remove excess adapter dimers from the library. All library construction steps were automated on the Revvity Janus platform. QPCR was performed using the Bio-Rad CFX384 Real-Time System and the KAPA SYBR FAST qPCR Master Mix (079559397001) in order to determine the nanomolarity of the final libraries. Sequencing was performed using the AVITI instrument and the Element Cloudbreak Freestyle High Output 300 cycle flow cell. Native Element Elevate libraries were constructed using the Elevate Mechanical Library Prep Kit (830-00008) for subsequent sequencing using the Element UltraQ sequencing kit. Samples were sheared on the Covaris R230 focused ultra-sonicator to a mean fragment size of 380bp. This was followed by a combined End-repair and A-tailing step and the subsequent ligation of full length Elevate adapters. Final libraries were subjected to a single Takara NucleoMag cleanup followed by a two-sided size selection NucleoMag cleanup. QPCR was performed using the Bio-Rad CFX384 Real-Time System and the KAPA SYBR FAST qPCR Master Mix (079559397001) in order to determine the nanomolarity of the final libraries. Sequencing was carried out on the AVITI instrument and the Element Cloudbreak UltraQ flow cell following the manufacturer’s instructions.

### Ultima UG100 and ppmSeq

PCR-free WGS libraries were generated from 500 ng of genomic DNA following the UG PCR-Free WGS Library Preparation Protocol for Solaris Free (document D1001056, revision 01). gDNA was enzymatically fragmented, end-repaired, and A-tailed using the NEBNext*®* Ultra II FS DNA Library Prep Kit (E7430). xGEN™ PCR-free Adapters for Ultima Genomics (Integrated DNA Technologies) were added via ligation and the resulting library molecules were size selected using a double-sided SPRI to capture average insert sizes of approximately 350-400 bp. ppmSeq*®* libraries were generated from 500 ng of genomic DNA following the UG ppmSeq Genomic DNA Library Preparation Protocol for Solaris Free (document D1000993, revision 02). gDNA was enzymatically fragmented using NEBNext*®* UltraShear (M7634) and end-repaired and A-tailed using the NEBNext*®* Ultra™ II End Repair/dA-Tailing Module (E7546). UG ppmSeq Adapters were added via ligation using the NEBNext*®* Ultra™ II Ligation Module (E7595). Resulting library molecules were size selected using a double-sided SPRI to capture average insert sizes of approximately 350-450 bp. Library size was determined via High Sensitivity D1000 TapeStation (Agilent Technologies), and the library concentration was quantified via QuantStudio qPCR (Thermo Fisher Scientific) using the NEBNext Library Quant Kit for Ultima Genomics (E3410). Libraries were pooled and diluted to a final concentration of 55pM in 1000μL.

### PacBio Onso

Onso libraries were prepared using the Onso Fragmentation DNA Library Preparation Kit according to the protocol “Preparing Onso Libraries from Genomic DNA for Short-Read Sequencing.” Libraries were quantified using the Onso Library Quantification Kit and subsequently sequenced on Onso flow cells with a 300-cycle kit (2x150bp) for 48 hours.

### MGI T7

Libraries were prepared using DNBSEQ Fast PCR-Free Library Prep set with 400ng input. Fragmentation time was 8 minutes, followed by double-sided size selection to select an average ∼475bp peak. dsDNA was denatured into single strands. The 5’ ends were phosphorylated and complimentary splint oligos were utilized to circularize the material into ssCircDNA. Remaining linear DNA was digested and 130 fmol ssCircDNA input was made into DNA nanoballs (DNBs) using the OneStep DNB Make kit. DNBs were loaded onto Complete Genomics flow cells and sequencing was performed on the DNBSEQ-T7 with a PE read length of 150bp

### Roche SBX-D

Libraries were prepared using the SBX-D SOP Version LP.11 (Roche Sequencing Solutions, Seattle, WA). Genomic DNA (50 ng) was diluted in 35 μL of 10 mM Tris-HCl, pH 8, followed by an addition of 25 μL fragmentation mix targeting a median insert size of >200bp. Fragmented DNA was purified using KAPA HyperPure beads at a 1.6X ratio. Fragmented DNA was subjected to an end-repair and A-tailing incubation. This was followed by ligation using Hairpin Sample Identification Adapter and Y-adapter. Ligated libraries were purified using KAPA HyperPure beads at a 0.33X ratio. Libraries underwent linear amplification for 6 hours and were then purified using KAPA HyperPure beads at a 0.89X ratio. Libraries were quantified on a Qubit^TM^ 4 Fluorometer (Thermo Fisher Scientific, Waltham, MA) using a custom-built assay (Roche Sequencing Solutions, Seattle, WA). Three picomoles of the prepared SBX-D library were used to synthesize and sequence Xpandomer molecules using prototype Axelios systems.

### Construction of the COLO829B-DSA

A high-quality diploid genome assembly of the COLO829BL lymphoblastoid cell line was assembled as part of the SMaHT Network using PacBio HiFi Fiber-seq, ultra-long Oxford Nanopore (ONT), and Hi-C data [24]. Problematic regions identified by *Flagger* [34] and *nucFreq* [35] were removed from consideration in the COLO829BL-DSA mapping cases. Collapsed and duplicated regions were also largely confined to centromeric and acrocentric regions, and these problematic intervals were masked before using the COLO829BL-DSA to directly quantify platform-specific SNMRs and IDMRs.

### Subsampling of data

Data reduction, or “downsampling,” was performed with *sambamba* (v0.6.8) [36] on the production-level FASTQ files to a range of fractional levels using the following command: *sambamba view -h -s [FRACTION] -f bam -o [OUTPUT_BAM_FILE] [INPUT_BAM_FILE]* (where -h means include the input BAM file header in the output BAM, -f indicates the format of the output and -o is the name of the output file).

### Mapping of sequencing reads

For the COLO829BL sample, we used the COLO829BL-DSA, a custom single-diplotype reference FASTA [24]. For the HG002 samples, we used the HG002-DSA; a single-diplotype reference FASTA [27]. For both cases of mapping to the respective DSAs and mapping to the generic GRCh38 human reference, reads generated from each of the sequencing platforms were aligned to the reference using *bwa-mem2* (v2.2.1) [37] with the following command: *bwa-mem2 mem [REFERENCE] [READ_1_FASTQ] [READ_2_FASTQ] > [OUTPUT_SAM_FILE]*.

### Creating and mapping DSA-derived simulated reads for baseline variation assessment

To estimate the contribution of genuine genomic variation independently of sequencing-induced errors, we simulated an idealized set of 151-bp paired-end reads from the COLO829BL-DSA reference. To make the simulated data comparable to the observed short-read datasets, we used the empirical MGI T7 COLO829BL run as a template for read length and insert-size distribution. We generated in silico read pairs from the COLO829BL-DSA FASTA sequence at the same mapping start positions and with the same insert sizes as the observed reads, while restricting simulation to genomic intervals accessible to short-read sequencing. The resulting reads were then aligned to GRCh38 under the same conditions used for the experimental datasets.

### Mismatch and base-quality recalibration analysis

Full, base-quality-independent SNMR and IDMR were computed from the raw alignment data and independently verified with *alfred* v0.5.1 [38]. Base-quality recalibration was performed with *best* v0.1.0 [39]by stratifying single-nucleotide mismatch observations into bins defined by the reported base-quality score and estimating the empirical relationship between mismatch frequency and base-quality value. The resulting recalibrated base qualities were then used to apply the BQ ≥ 30 threshold for downstream filtered mismatch analyses, thereby restricting comparisons to a standardized high-confidence subset. All tools were run on BAM files generated by *bwa-mem2*.

### High-quality allele mismatch calculation

For SNMR calculation, BAM alignments were required to include both a CIGAR string [40] and a “matching” (MD) string. The CIGAR string was used to identify the number and genomic position of insertions and deletions, whereas the MD string provided the location of single-nucleotide mismatches and the deleted reference sequence. This approach was first validated against the *alfred* (v0.5.1) and then adapted to our analysis framework. Relative to *alfred*, our implementation reproduced SNMR values within <0.3% and IDMR values within <3%; in both cases, *alfred* slightly underestimated mismatch counts.

For insertions, the quality score was defined as the mean base quality of all nucleotides in the inserted segment. High-quality deletions were not analyzed because deleted bases do not have associated quality scores in BAM files, making a consistent deletion-quality estimate impossible. Although the CIGAR string left-justifies inserted homopolymer bases relative to the reference, base qualities are typically assigned to inserted alleles using right-justified coordinates. Because some insertion qualities appeared to be left-justified in certain cases, we conservatively took the minimum quality between the left- and right-justified assignments to avoid overestimating the number of high-quality mismatched nucleotides.

### Low-complexity region determination and analysis

We started with the Global Alliance for Genomics and Health’s (GA4GH)-defined low-complexity regions (LCRs) [17], which include tandem repeats and homopolymer regions in GRCh38, and lifted these regions over to the COLO829BL-DSA using *rustybam* [41]. For this analysis, BQ ≥ 30 single-nucleotide and insertion mismatches within LCRs and non-LCRs were normalized to the number of sequenced bases in each region class and to the relative genomic size of the corresponding region set. In addition, rates were adjusted for the fraction of BQ ≥ 30 bases in each platform’s full dataset.

## Declarations

### Ethics approval and consent to participate

’Not applicable’

### Consent for publication

’Not applicable’

### Availability of data and materials

This study was conducted as part of the NIH Common Fund consortium initiative, Somatic Mosaicism across Human Tissues (SMaHT). More information about the SMaHT Network and resources can be found online [42].

Analysis scripts used to create and summarize results have been deposited in GitHub and are available publicly [43].

Any additional information required to reanalyze the data reported in this work is available from the lead contact upon request.

The benchmark datasets described in this study are currently available through the SMaHT Data Portal [44] and the data is accessible to anyone after the creation of an account.

## Authors’ Contributions

S.R.M., J.D.S., A.B.S, E.E.E and C.L.W. conceptualized the project. S.R.M. and M.R.V. performed tool development. S.R.M. and Y.K. conducted bioinformatics analyses. S.R.M., J.D.S., C.D.F, and C.L.W. wrote the manuscript with input from all authors. C.L.W. supervised the project. All authors reviewed and approved the final manuscript.

## Acknowledgements

The authors want to acknowledge Colleen Davis, Jeffrey Ou, Karynne Patterson and Christina Zakarian for support in preparing the manuscript. Brian Coullahan and Semyon Kruglyak from Element Biosciences; Noëlle Bittner, Siyuan Zhang, and Ian McLaughlin at Pacific Biosciences; Abel Gutierrez, Nidhanjali Bansal, Biniam Feleke and Corey Jefferson at Complete Genomics; Alix Cruse, Ariel Jaimovich, Gat Krieger, Itai Rusinek and Ilya Soifer at Ultima Genomics; and Jagdeesh Chandrasekar, Mahdi Golkaram, Chen Zhao, Alberto Gatto, Cynthia Cech, Kendall Berg, John Mannion and Mark Kokoris from Roche Sequencing Solutions.

## Supplemental Information

**Additional file 1**: Overview of the technical features and sequencing chemistries for the nine short-read sequencing platforms evaluated in this analysis.

## Supplementary Tables

Table S1. Key features of the short-read sequencing platforms used in this study.

Table S2. Consistency of mismatch rates across a range of depths of coverage generated by downsampling the same set of FASTQs to different fractional levels.

Table S3. SNMRs for all platforms and each sample mapped to GRCh38 and its DSA. Table S4. IDMRs for all platforms and each sample mapped to GRCh38 and its DSA.

Table S5. SNMRs and DMRs for only the highest-quality events (BQ ≥ 30) for all platforms and each sample mapped to its respective DSA.

Table S6. Proportion of mismatch errors associated with each reference base and their altered sequences.

Table S7. Genomic complexity effects on SNMRs and DMRs for high-confidence events (BQ ≥ 30) across platforms and samples.

