## Supplemental Materials for "Short-Read Sequencing Benchmarking with Donor-Specific Assemblies"

**Supplementary Information**

Overview of the technical features and sequencing chemistries for the nine short-read sequencing platforms evaluated in this analysis.

**Supplementary Tables:**

Table S1. Key features of the short-read sequencing platforms used in this study.

Table S2. Consistency of mismatch rates across a range of depths of coverage generated by downsampling the same set of FASTQs to different fractional levels.

Table S3. SNMRs for all platforms and each sample mapped to GRCh38 and its DSA.

Table S4. IDMRs for all platforms and each sample mapped to GRCh38 and its DSA.

Table S5. SNMRs and DMRs for only the highest-quality events (BQ ≥ 30) for all platforms and each sample mapped to its respective DSA.

Table S6. Proportion of mismatch errors associated with each reference base and their altered sequences.

Table S7. Genomic complexity effects on SNMRs and DMRs for high-confidence events (BQ ≥ 30) across platforms and samples.

### 1. Illumina NovaSeq

Illumina's sequencing-by-synthesis (SBS) technology is the most broadly adopted short-read sequencing technology. Launched in 2006, this technology is used on the NovaSeq 6000 and NovaSeq X Plus instruments. Library molecules are bound to a lawn of oligos as they cross the flow cell surface. Bridge amplification generates additional clonal molecules and forms clusters, which are then detected by the instrument's optics. SBS functions by flooding the flow cell with fluorescently labeled bases. Complementary bases bind to the library of molecules and blocking prevents multiple bases from being incorporated simultaneously. Lasers excite the fluorophore, which indicates which base was incorporated. Prior to advancing the next cycle, the fluorophore and block must be removed. This process results in molecular scarring, which can interfere with subsequent nucleotide addition. Illumina flow cells are patterned, meaning that nanowells are etched on the surface of the flow cell in a regular pattern. Clusters can only form in nanowells, which allows for more efficient use of the flow cell surface and faster imaging, since cluster positions are known. Shorter library molecules migrate across the flow cell surface faster than larger molecules, leading to a bias towards smaller molecules. This is more pronounced on patterned flow cells than on unpatterned flow cells. Illumina NovaSeq X instruments use a new form of SBS chemistry called XLEAP-SBS, which results in faster cycle times and higher fidelity. The NovaSeq X instrument can generate more than 7 trillion bases per run. The flow cell consists of 8 individually addressable lanes, reducing the need for a lab to maintain a high number of index combinations.

### 2. Element AVITI

Element Biosciences's AVITI platform uses avidity base chemistry (ABC), which they have named Cloudbreak, for sequencing. Their Cloudbreak freestyle version of this ABC chemistry allows the user to load premade libraries containing Illumina or Element-specific adapters. Linear library molecules are circularized on the flow cell and then rolling circle amplification generates clonal clusters, termed 'polonies'. The polony sequence is determined by introducing fluorescently labeled base-specific avidites. When the polymerase adds the base the avidite is held by the polony. The fluorophore is then excited, and the polony is imaged. The avidite is removed and the next cycle begins. Avidite sequencing has the benefit of not requiring the removal of a blocker, which minimizes DNA damage concerns. Reagent costs are potentially lower because fluorophores are bound to the avidite and not the individual nucleotides. Element flow cells are unpatterned and consist of two individually addressable lanes. The instrument is capable of being run in what Element terms 'High Density' mode, which generates more data than the standard run mode. This is achieved by relaxing some of the polony filtering. Element Biosciences UltraQ technology uses Cloudbreak sequencing with a few modifications to generate data with a higher average quality score. Native Element library molecules are loaded onto the instrument and treated with a deamination reagent before polony generation, which digests molecules with deamination damage. This results in the removal of molecules that have the potential to introduce errors downstream. Second, Element introduces dark cycling at the beginning of read two in order to skip over bases that have a higher error rate due to the end repair process during library construction. In February 2026, Element announced the Vitari instrument, which is a higher throughput platform claiming up to 10 billion reads per 6 lane flow cell, using the same chemistry as the AVITI.

**3. MGI T7**

MGI sequencing technology uses rolling circle amplification to generate what MGI calls 'DNA Nanoballs'. Like Element AVITI sequencing, the use of rolling circle amplification reduces the likelihood of PCR errors being perpetuated. Nanoballs are introduced to the patterned flow cell where they bind to the positively charged surface. MGI sequencing technology leverages combinatorial probe-anchor synthesis (cPAS) to interrogate the library sequence using four fluorescently labeled probes. These probes feature reversible terminators and are added during sequential cycles by a DNA polymerase. Lasers excite the fluorescent labels and the instrument registers the base. For the analysis presented here, data was generated on the high throughput DNBSEQ-T7 instrument. In the fall of 2025, MGI announced the T7+ instrument, which promises more than double the output per flow cell with higher average Q-scores.

**4. PacBio Onso**

Pacific Biosciences Onso short-read sequencing platform uses sequencing by binding (SBB) technology, which separates the interrogation and integration steps to reduce sequencing errors. In the first step, a block is applied to the strand of nucleotides, allowing only one fluorescently labeled base to be added in step 2. The flow cell is imaged, and the labeled nucleotide is removed. The block is then moved in step 3, allowing a base to be added in step 4. The process is then repeated. Prior to sequencing, libraries are loaded onto the flowcell and cluster generation occurs on a separate instrument. This instrument has the highest cost and one of the longest run times of the sequencing platforms considered here.  In January 2026, the technology underlying the Onso platform was sold to Illumina and is not expected to be marketed in its current form.

### 5. Ultima UG100

Ultima Genomics has developed a sequencing technology that uses a wafer from the semiconductor industry as a flow cell. These readily available discs reduce costs relative to the proprietary flow cells used by other technologies. Inside the UG100 instrument the disc spins like a CD or DVD and reagents are applied to the surface sequentially. Optics are fixed and read the disc as it spins. This is the opposite of other sequencing technologies where the optical module moves over the fixed flow cell. Ultima uses a flow chemistry that results in single end sequencing. A mix of natural and labeled nucleotides flow across the wafer surface one base at a time and are incorporated into the growing strand. Multiple nucleotides can be incorporated in a single flow in homopolymer regions. Fluorescence intensity is used to determine the number of bases added. The use of unlabeled nucleotides reduces the cost of reagents. The UG100 has large footprint and requires emulsion PCR to clonally amplify the sequencing libraries. Loading times are flexible, allowing for continuous operation and high throughput data generation.  Unlike a traditional flow cell, Ultima's spinning wafer technology does not have lane compartment, therefore, it requires a significant number of indexes to pool multiple libraries for highly multiplexed sequencing runs to take full advantage of the large data generation capacity.

Ultima Genomics ppmSeq captures double stranded DNA fragments onto beads that are clonally amplified during emulsion PCR. Contained in the adapter portion of the library molecule is a string of adenine bases. Post amplification each bead should contain a mix of A and T in this region. ppmSeq then uses standard Ultima sequencing methods on the UG100 to sequence the duplex reads. During read processing, the mix of A and T bases in the adapter is used to ensure that both strands of the duplex molecule are equally represented. Reads where there is a variant on only the forward or reverse strand are filtered out, leaving only concordant reads between the two strands. Because of this read filtering, ppmSeq requires greater sequencing depth. Q-scores for ppmSeq can exceed 60, resulting in higher confidence in detecting rare variants. In early 2026, Ultima announced the UG200 instrument with a claimed output twice the UG100. UG200 workflow uses isothermal amplification instead of emulsion PCR for template amplification, which eliminates the need for the dedicated emulsion PCR instrument.

### 6. Roche Axelios 1

Roche's Sequencing-by-Expansion (SBX) is the latest short-read sequencing technology released commercially in mid-2026. SBX technology is being developed by Roche after acquiring Stratos Genomics and Genia Technologies in 2020 and 2014, respectively. SBX uses nanopore-based sequencing technology. Molecules are detected as they pass through a nanopore and the detection signal is subsequently converted into a base call. Prior to nanopore sequencing, the DNA template strand is copied using synthetic expandable bases, which when incorporating into the DNA strand, expand significantly and form longer molecules. Roche refers to this molecule as an Xpandomer. The synthetic polymer is fed through the nanopore and the larger molecules result in a higher signal to noise ratio. One of the advantages of this technology is the high throughput where more than 5B duplex reads or seven 30x whole genomes can be sequenced in an hour. SBX-D is a duplex variation of this process where a hairpin adapter is substituted at one end of the molecule. Like ppmSeq both strands are read during the sequencing reaction and molecules where the duplex reads disagree are discarded. The system consists of two instruments: one for Xpandomer formation and the second for nanopore sequencing. Reagent costs were announced in February 2026 with the cost of a 30x human genome at $150 using the SBX-D workflow based on reusing the sensor module up to twenty times.

**Table S1. Key features of the short-read sequencing platforms used in this study**

| Technology | NovaSeq X | NovaSeq 6000 | UG100 | ppmSeq | T7 | AVITI | UltraQ | AXELIOS 1 SBX-D | Onso |
|---|---|---|---|---|---|---|---|---|---|
| **Manufacturer** | Illumina | Illumina | Ultima | Ultima | MGI | Element | Element | Roche | PacBio |
| **Read Structure** | PE | PE | SE | Duplex | PE | PE | PE | Duplex | PE |
| **Amplification** | Bridge Amplification | Bridge Amplification | Emulsion PCR | Emulsion PCR | Rolling Circle | Rolling Circle | Rolling Circle | Linear | Undisclosed |
| **Flow Cell Configuration** | 25B | S4 | Wafer | Wafer | High Throughput V3 | High Output | UltraQ High Output | Nanopore Sensor Module | - |
| **Run Time (300 cycles)**** | 48h | 44hr | 14h | 14hr | 24h | 38h | 44hr | 4hrs | 48h |
| **Reads (B)** | 52 | 20 | 8 | 8 | 5.8 | 1 | 0.8 | 5.3 | 1 |
| **Output (GB)** | 8000 | 3000 | 2400 | | 1750 | 300 | 240 | 2,300 | 150 |
| **Quality (Q30)** | >85% | | >85% | | >85% | >90% | >90% Q40 | Q39 Avg | >90% Q40 |
| **Cost/GB** | $2 | $5.50 | $1 | | $1.50 | $5.60 | $11.16 | $1.04 | $15 |

*Output is calculated per flow cell. All values are based on specifications published by the manufacturer.
**SBX sequencing does not involve cycling

**Table S2.** Consistency of mismatch rates across a range of depths of coverage generated by downsampling the same set of FASTQs to different fractional levels

| Platform | Sample | Reference | Gbs | SNMR | IDMR (insertions) | IDMR (deletions) |
|---|---|---|---|---|---|---|
| **Ultima (UG100)** | **HG002** | **GRCh38** | 3.4 | 4.3229 | 2.0303 | 1.8988 |
| | | | 13.6 | 4.3218 | 2.0300 | 1.8989 |
| | | | 27.1 | 4.3212 | 2.0297 | 1.8988 |
| | | | 54.3 | 4.3207 | 2.0294 | 1.8984 |
| | | | **std dev** | **9.28E-04** | **3.89E-04** | **2.06E-04** |

**Table S3.** SNMRs for all platforms and each sample mapped to GRCh38 and its DSA.

| Platform | Sample | Reference | SNMR |
|---|---|---|---|
| **Illumina (NovaSeq 6000)** | COLO829BL | GRCh38 | 5.50 |
| | | COLO829BL-DSA | 3.57 |
| | HG002 | GRCh38 | 5.60 |
| | | HG002-DSA | 3.51 |
| **Illumina (NovaSeq X)** | COLO829BL | GRCh38 | 4.00 |
| | | COLO829BL-DSA | 1.87 |
| | HG002 | GRCh38 | 4.23 |
| | | HG002-DSA | 2.14 |
| **Element (AVITI)** | COLO829BL | GRCh38 | 5.17 |
| | | COLO829BL-DSA | 3.20 |

| | | | |
|---|---|---|---|
| | HG002 | GRCh38 | 4.52 |
| | | HG002-DSA | 2.41 |
| **Element (AVITI w/UltraQ)** | COLO829BL | GRCh38 | 3.09 |
| | | COLO829BL-DSA | 0.98 |
| | HG002 | GRCh38 | 3.19 |
| | | HG002-DSA | 0.97 |
| **PacBio (Onso)** | COLO829BL | GRCh38 | 2.70 |
| | | COLO829BL-DSA | 0.49 |
| | HG002 | GRCh38 | 2.86 |
| | | HG002-DSA | 0.61 |
| **MGI (T7)** | COLO829BL | GRCh38 | 3.38 |
| | | COLO829BL-DSA | 1.61 |
| | HG002 | GRCh38 | 3.50 |
| | | HG002-DSA | 1.59 |
| **Ultima (UG100)** | COLO829BL | GRCh38 | 3.89 |
| | | COLO829BL-DSA | 1.80 |
| | HG002 | GRCh38 | 4.32 |
| | | HG002-DSA | 2.05 |
| **Ultima (ppmSeq)** | COLO829BL | GRCh38 | 2.84 |
| | | COLO829BL-DSA | 0.92 |
| | HG002 | GRCh38 | 2.80 |
| | | HG002-DSA | 0.59 |
| **151-bp reads generated from COLO829BL-DSA** | COLO829BL | GRCh38 | 1.57 |
| | | COLO829BL-DSA | 0.00 |

**Table S4.** IDMRs for all platforms and each sample mapped to GRCh38 and its DSA.

| Platform | Sample | Reference | IDMR x $10^{-3}$ (insertions) | IDMR x $10^{-3}$ (deletions) |
|---|---|---|---|---|
| **Illumina (NovaSeq 6000)** | COLO829BL | GRCh38 | 117.54 | 114.02 |
| | | COLO829BL-DSA | 9.11 | 8.90 |
| | HG002 | GRCh38 | 124.84 | 123.00 |
| | | HG002-DSA | 9.24 | 11.89 |

| | | | | |
|---|---|---|---|---|
| **Illumina (NovaSeq X)** | COLO829BL | GRCh38 | 126.03 | 119.09 |
| | | COLO829BL-DSA | 7.26 | 7.61 |
| | HG002 | GRCh38 | 126.41 | 122.58 |
| | | HG002-DSA | 8.21 | 8.55 |
| **Element (AVITI)** | COLO829BL | GRCh38 | 115.72 | 111.01 |
| | | COLO829BL-DSA | 4.32 | 3.52 |
| | HG002 | GRCh38 | 124.58 | 117.37 |
| | | HG002-DSA | 3.93 | 3.02 |
| **Element (AVITI w/UltraQ)** | COLO829BL | GRCh38 | 116.84 | 109.26 |
| | | COLO829BL-DSA | 2.95 | 2.34 |
| | HG002 | GRCh38 | 118.73 | 112.22 |
| | | HG002-DSA | 2.78 | 2.45 |
| **PacBio (Onso)** | COLO829BL | GRCh38 | 132.20 | 122.94 |
| | | COLO829BL-DSA | 5.22 | 5.34 |
| | HG002 | GRCh38 | 127.02 | 115.01 |
| | | HG002-DSA | 7.14 | 3.08 |
| **MGI (T7)** | COLO829BL | GRCh38 | 102.42 | 112.64 |
| | | COLO829BL-DSA | 4.61 | 14.77 |
| | HG002 | GRCh38 | 101.46 | 103.34 |
| | | HG002-DSA | 3.11 | 4.08 |
| **Ultima (UG100)** | COLO829BL | GRCh38 | 1944.23 | 2019.85 |
| | | COLO829BL-DSA | 1816.64 | 1913.88 |
| | HG002 | GRCh38 | 2029.86 | 1899.09 |
| | | HG002-DSA | 1928.42 | 1816.13 |
| **Ultima (ppmSeq)** | COLO829BL | GRCh38 | 1522.14 | 1214.69 |
| | | COLO829BL-DSA | 1420.48 | 1124.24 |
| | HG002 | GRCh38 | 1158.08 | 946.79 |
| | | HG002-DSA | 1055.33 | 857.60 |
| **151-bp reads generated from COLO829BL-DSA** | COLO829BL | GRCh38 | 93.70 | 91.18 |
| | | COLO829BL-DSA | 0.00 | 0.00 |

**Table S5.** SNMRs and IDMRs for only the highest-quality events (BQ ≥ 30) for all platforms and each sample mapped to its respective DSA

| Platform | Sample | Reference | SNMR x $10^{-3}$ | IDMR x $10^{-3}$ (insertions) |
|---|---|---|---|---|
| **Illumina (NovaSeq 6000)** | COLO829BL | COLO829BL-DSA | 185.05 | 3.21 |
| | HG002 | HG002-DSA | 180.96 | 4.41 |
| **Illumina (NovaSeq X)** | COLO829BL | COLO829BL-DSA | 202.47 | 4.41 |
| | HG002 | HG002-DSA | 151.98 | 5.13 |
| **Element (AVITI)** | COLO829BL | COLO829BL-DSA | 126.07 | 2.64 |
| | HG002 | HG002-DSA | 48.16 | 1.93 |
| **Element (AVITI w/UltraQ)** | COLO829BL | COLO829BL-DSA | 86.96 | 2.37 |
| | HG002 | HG002-DSA | 34.91 | 1.65 |
| **PacBio (Onso)** | COLO829BL | COLO829BL-DSA | 144.21 | 4.88 |
| | HG002 | HG002-DSA | 57.94 | 5.26 |
| **MGI (T7)** | COLO829BL | COLO829BL-DSA | 188.43 | 4.11 |
| | HG002 | HG002-DSA | 80.03 | 2.67 |
| **Ultima (UG100)** | COLO829BL | COLO829BL-DSA | 306.81 | 32.03 |
| | HG002 | HG002-DSA | 350.81 | 37.20 |
| **Ultima (ppmSeq)** | COLO829BL | COLO829BL-DSA | 167.73 | 28.23 |
| | HG002 | HG002-DSA | 95.74 | 38.30 |
| **Roche (SBX-D)** | COLO829BL | COLO829BL-DSA | 95.00 | 30.54 |
| | HG002 | HG002-DSA | 24.07 | 30.95 |

**Table S6.** Proportion of mismatch errors associated with each reference base and their altered sequences

| HG002 | | | | | | | | | |
|---|---|---|---|---|---|---|---|---|---|
| **Ref>Alt** | **Illumna 6000** | **Novoseq X plus** | **Element Aviti** | **Element UltraQ** | **PacBio Onso** | **MGI T7** | **Ultima UG-100** | **Ultima ppmseq** | **Roche SBX-D** |
| A>T | 5.9% | 3.2% | 7.9% | 11.8% | 6.5% | 15.6% | 10.4% | 10.6% | 7.2% |
| A>C | 10.0% | 8.7% | 11.6% | 7.8% | 4.2% | 4.4% | 8.7% | 8.2% | 4.6% |
| A>G | 13.2% | 13.5% | 3.3% | 3.0% | 13.0% | 6.3% | 13.4% | 11.4% | 15.2% |
| T>A | 6.1% | 3.2% | 8.0% | 12.0% | 6.5% | 15.6% | 10.6% | 10.8% | 7.3% |
| T>C | 13.7% | 14.2% | 4.9% | 4.8% | 15.0% | 7.2% | 13.6% | 12.0% | 18.8% |
| T>G | 10.2% | 8.8% | 11.8% | 8.0% | 4.3% | 4.5% | 8.6% | 8.0% | 5.1% |
| C>A | 10.5% | 9.9% | 11.1% | 11.9% | 4.9% | 2.9% | 5.3% | 7.1% | 4.4% |
| C>T | 6.6% | 10.7% | 11.9% | 11.6% | 16.5% | 9.8% | 6.0% | 8.4% | 13.9% |
| C>G | 3.6% | 4.0% | 3.8% | 3.3% | 4.5% | 10.9% | 6.1% | 4.2% | 3.8% |
| G>A | 6.2% | 10.0% | 10.7% | 10.3% | 15.1% | 9.2% | 5.8% | 7.7% | 11.3% |
| G>T | 10.5% | 9.9% | 11.2% | 12.2% | 5.1% | 2.9% | 5.4% | 7.4% | 4.7% |
| G>C | 3.6% | 4.0% | 3.8% | 3.3% | 4.5% | 10.9% | 6.0% | 4.1% | 3.7% |
| **COLO829BL** | | | | | | | | | |
| **Ref>Alt** | **Illumna 6000** | **Novoseq X plus** | **Element Aviti** | **Element UltraQ** | **PacBio Onso** | **MGI T7** | **Ultima UG-100** | **Ultima ppmseq** | **Roche SBX-D** |
| A>T | 4.3% | 2.1% | 5.9% | 5.8% | 11.6% | 11.4% | 10.1% | 7.4% | 3.6% |
| A>C | 5.0% | 6.5% | 8.0% | 4.2% | 2.6% | 1.0% | 7.4% | 5.1% | 3.1% |
| A>G | 15.8% | 16.2% | 12.7% | 14.8% | 12.2% | 7.6% | 13.4% | 14.7% | 20.7% |
| T>A | 4.4% | 2.2% | 6.0% | 6.0% | 11.4% | 11.5% | 10.4% | 7.6% | 3.8% |
| T>C | 17.4% | 17.6% | 13.7% | 17.4% | 13.2% | 9.0% | 15.0% | 16.4% | 25.0% |
| T>G | 5.2% | 6.7% | 8.4% | 4.7% | 3.1% | 1.2% | 7.2% | 5.2% | 3.1% |
| C>A | 8.7% | 10.6% | 7.6% | 6.9% | 6.5% | 3.0% | 5.3% | 7.3% | 2.9% |
| C>T | 8.1% | 9.9% | 11.5% | 13.9% | 14.0% | 12.6% | 8.6% | 11.8% | 15.4% |
| C>G | 7.2% | 3.8% | 3.2% | 2.4% | 2.4% | 13.4% | 4.4% | 2.6% | 2.0% |
| G>A | 8.2% | 10.0% | 12.0% | 14.6% | 14.1% | 12.8% | 8.5% | 11.8% | 15.3% |
| G>T | 8.6% | 10.5% | 7.7% | 7.0% | 6.6% | 3.0% | 5.5% | 7.5% | 2.8% |
| G>C | 7.3% | 4.1% | 3.3% | 2.5% | 2.5% | 13.4% | 4.3% | 2.6% | 2.3% |

**Table S7**. Genomic complexity effects on SNMRs and DMRs for high-confidence events (BQ ≥ 30) across platforms and samples.

| Platform | Sample | Reference | Region[a] | SNMR x $10^{-3}$ | IDMR x $10^{-3}$ (insertions) |
|---|---|---|---|---|---|
| **Illumina (NovaSeq 6000)** | COLO829BL | COLO829BL-DSA | LCR | 539.21 | 113.44 |
| | | | non-LCR | 32.63 | 0.16 |
| | HG002 | HG002-DSA | LCR | 611.95 | 69.44 |
| | | | non-LCR | 34.78 | 0.71 |
| **Illumina (NovaSeq X)** | COLO829BL | COLO829BL-DSA | LCR | 561.44 | 132.25 |
| | | | non-LCR | 33.93 | 0.21 |
| | HG002 | HG002-DSA | LCR | 614.37 | 70.28 |
| | | | non-LCR | 26.91 | 0.74 |
| **Element (AVITI)** | COLO829BL | COLO829BL-DSA | LCR | 255.48 | 70.86 |
| | | | non-LCR | 22.09 | 0.10 |
| | HG002 | HG002-DSA | LCR | 133.22 | 8.18 |
| | | | non-LCR | 9.07 | 0.35 |
| **Element (AVITI w/UltraQ)** | COLO829BL | COLO829BL-DSA | LCR | 125.79 | 59.59 |
| | | | non-LCR | 13.51 | 0.07 |
| | HG002 | HG002-DSA | LCR | 98.22 | 6.45 |
| | | | non-LCR | 6.29 | 0.29 |
| **PacBio (Onso)** | COLO829BL | COLO829BL-DSA | LCR | 385.78 | 101.45 |
| | | | non-LCR | 21.23 | 0.25 |
| | HG002 | HG002-DSA | LCR | 173.28 | 12.08 |

| | | | | | |
|---|---|---|---|---|---|
| | | | non-LCR | 9.98 | 0.94 |
| **MGI (T7)** | COLO829BL | COLO829BL-DSA | LCR | 336.45 | 108.79 |
| | | | non-LCR | 34.19 | 0.16 |
| | HG002 | HG002-DSA | LCR | 218.89 | 9.13 |
| | | | non-LCR | 14.89 | 0.49 |
| **Ultima (UG100)** | COLO829BL | COLO829BL-DSA | LCR | 773.80 | 634.54 |
| | | | non-LCR | 63.44 | 51.62 |
| | HG002 | HG002-DSA | LCR | 1108.38 | 679.29 |
| | | | non-LCR | 74.67 | 57.78 |
| **Ultima (ppmSeq)** | COLO829BL | COLO829BL-DSA | LCR | 358.52 | 75.50 |
| | | | non-LCR | 31.39 | 7.33 |
| | HG002 | HG002-DSA | LCR | 426.36 | 94.71 |
| | | | non-LCR | 17.41 | 9.27 |
| **Roche (SBX-D)** | COLO829BL | COLO829BL-DSA | LCR | 141.06 | 262.33 |
| | | | non-LCR | 14.54 | 6.61 |
| | HG002 | HG002-DSA | LCR | 123.46 | 212.35 |
| | | | non-LCR | 4.02 | 6.97 |

**a** Low-complexity regions (LCRs) are defined by the GA4GH consortium tandem repeat and homopolymer regions in GRCh38 and then lifted-over to the COLO829BL DSA. Non-LCR refers to all regions of the genome outside of LCRs.
